# Change point estimation by the mouse medial frontal cortex during probabilistic reward learning

**DOI:** 10.1101/2022.05.26.493245

**Authors:** Cayla E. Murphy, Hongli Wang, Heather K. Ortega, Alex C. Kwan, H. Atilgan

## Abstract

There are often sudden changes in the state of environment. For a decision maker, accurate prediction and detection of change points are crucial for optimizing performance. Still unclear, however, is whether rodents are simply reactive to reinforcements, or if they can be proactive to estimate future change points during value-based decision making. In this study, we characterize head-fixed mice performing a two-armed bandit task with probabilistic reward reversals. Choice behavior deviates from classic reinforcement learning, but instead suggests a strategy involving belief updating, consistent with the anticipation of change points to exploit the task structure. Excitotoxic lesion and optogenetic inactivation implicate the anterior cingulate and premotor regions of medial frontal cortex. Specifically, over-estimation of hazard rate arises from imbalance across frontal hemispheres during the time window before the choice is made. Collectively, the results demonstrate that mice can capitalize on their knowledge of task regularities, and this estimation of future changes in the environment may be a main computational function of the rodent dorsal medial frontal cortex.

## Introduction

In life, we experience twists and turns – discrete events that abruptly alter the state of environment. In some cases, the change is a one-time occurrence that is impossible to predict. We must then adjust by assessing the new situation following the change. However, in other cases, the changes may occur repeatedly with certain tendencies. For example, a favorite chef in a restaurant may have a recurring schedule where she cooks throughout the year, except in the summer for 3 – 5 weeks when she would take a vacation and lets a substitute take over. As a patron, it would be advantageous to learn this pattern, anticipate the impending switches, and maximize the chance of receiving a delicious outcome. While it is evident that humans can estimate change points and leverage the information in decision-making, whether animals such as mice have this ability and the neural substrates supporting the computations remain unclear.

A classic paradigm to study decision-making in response to repeated changes is the two-armed bandit task. Each trial, the animal has two options, and each option is associated probabilistically with a reward. After a certain number of trials, the reward probabilities are switched among the options. The two-armed bandit task is widely used because it can be tested in different species including humans (Evers et al., 2005; O’Doherty et al., 2001; Tsuchida et al., 2010), monkeys (Clarke et al., 2008; Costa et al., 2015; Donahue and Lee, 2015; Samejima et al., 2005), rats (Groman et al., 2019; Hamid et al., 2016; Ito and Doya, 2009; Sul et al., 2011; Sul et al., 2010), and mice (Bari et al., 2019; Grossman et al., 2022; Hattori et al., 2019; Tai et al., 2012). Moreover, the paradigm has translational significance because it can reveal defects from pharmacological interventions (Costa et al., 2015) or in animal models for psychiatric disorders (Groman et al., 2018; Liao and Kwan, 2021; Villiamma et al., 2022).

Most analyses of rodents performing two-armed bandit and related decision-making tasks have relied on simple reinforcement learning schemes such as Q-learning algorithms (Bari et al., 2019; Groman et al., 2019; Hattori et al., 2019; Ito and Doya, 2009; Sul et al., 2010; Wang et al., 2022), with some exceptions (Beron et al., 2022; Ito and Doya, 2015). Q-learning algorithms assume that animals learn from experience, and therefore choice behavior adapts only after a change point has occurred. By contrast, recent studies in monkeys and humans have challenged this assumption.

Namely, primates can exploit predictable structure in a task and adjust for an impending change point (Bartolo and Averbeck, 2020; Costa et al., 2015; Jang et al., 2019; Woo et al., 2023). Indeed, under some situations, rodents also seem to make inferences about hidden states (Liu et al., 2021; Starkweather et al., 2017; Starkweather et al., 2018; Vertechi et al., 2020; Woo et al., 2023). These recent results hint at the possibility that mice may leverage their knowledge of task structure for probabilistic reward learning.

To test the possibility that rodents may estimate change points during probabilistic reward learning, we trained head-fixed mice on a two-armed bandit task. By analyzing a sizable data set totaling 1,007 sessions involving 15,352 reversals, we demonstrate that mice are sensitive to impending switches in reward probabilities, because they alter their choices prior to the actual change points. We show that the animals’ choice behavior can be modelled effectively with a Bayesian framework involving belief updating and choice kernels. Furthermore, we performed unilateral and bilateral excitotoxic lesions and optogenetic inactivation to demonstrate mechanistically how the anterior cingulate and premotor regions of medial frontal cortex may be involved in the computation. Together, the results indicate that mice can take advantage of the task structure to solve a classic probabilistic reward learning task and implicate the dorsal medial frontal cortex as a locus in the accurate estimation of future changes in the state of environment.

## Results

### Mice use their knowledge of the task structure during a two-armed bandit task

We trained head-fixed C57BL/6J mice on a two-armed bandit task involving probabilistic reward reversals. On each trial, the mouse could choose left or right by a directional tongue lick. The two options were associated with different reward probabilities, e.g., “70:10” for 70% and 10% chance to receive water from the left and right spouts respectively (**Figure 1A – B**). The reward probabilities would flip when the animal reaches the switching condition, which is a performance-dependent number of trials to fulfill a criterion (L_Criterion_, 10 trials choosing the better option) followed by a performance-independent random number of trials (L_Random_, drawn from a geometric distribution with *p* = 0.0909 and truncated at 30). In an example session shown in **Figure 1C**, the animal performed more than 500 trials, including 15 reversals of 70:10 and 10:70 blocks. To visualize how animals adjust to the sudden changes in reward probabilities, we aligned the trials by the time of block switches. As expected, mice primarily chose the better option pre-switch, and then quickly adapted their preferred action post-switch (**Figure 1D**, n = 31 mice, 617 sessions, 9,163 blocks).

**Figure 1:**
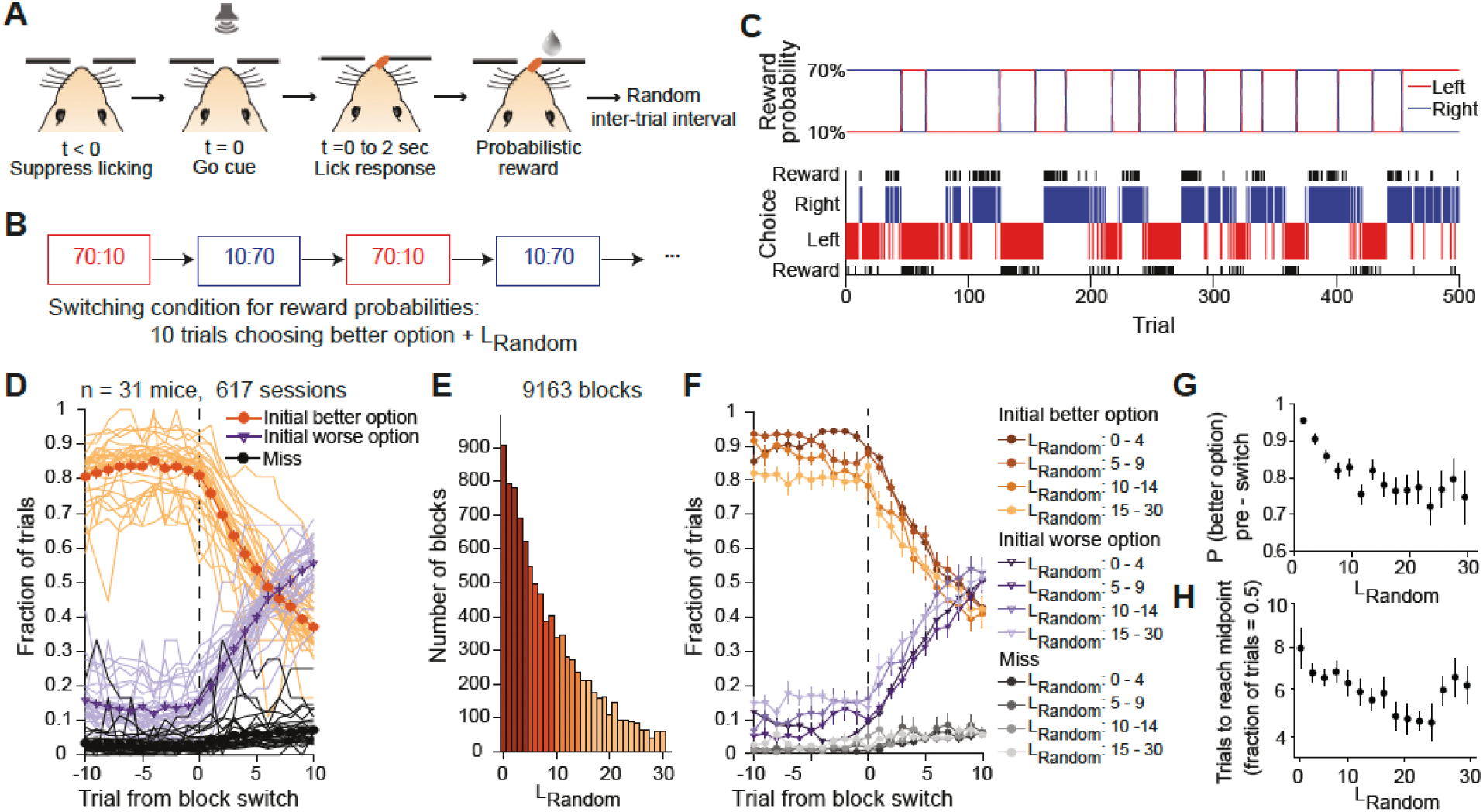
Mice were sensitive to block length and leverage this information during the two-armed bandit task. **(A)** The mouse makes a left or right choice via tongue lick after the go cue. Depending on the reward probabilities, the choice might lead to water. **(B)** Trials were organized into blocks, each with distinct reward probabilities: “70:10” (70% chance to receive water for left choice; 10% for right) or “10:70” (10% for left; 70% for right). The block switches after the animal choose the high-reward-probability side ten times (L_Criterion_) plus an additional random number of trials (L_Random_, drawn from exponential distribution, up to 30 trials). **(C)** Performance of a mouse in one example session. The top row shows reward probabilities for left and right options. The bottom row shows the animal’s choices and the outcomes. **(D)** Choice behavior around block switches. Thin line, mean values for individual animal. Thick line, mean values and SEM for all animals. **(E)** Histogram of L_Random._ For all blocks with L_Criterion_ ≤ 20. Colors indicate the 4 ranges of L_Random_ for subsequent analyses. **(F)** Choice behavior around block switches, plotted separately for the 4 ranges of L_Random_. Mean values and SEM for all animals. **(G)** The probability of choosing the better option on the trial immediately preceding the switch, as a function of L_Random_ for the block preceding the switch. Mean values and SEM for all animals. (H) The number of trials to reach midpoint (when animal is equally likely to choose either option) as a function of L_Random_ for the block preceding the switch. Mean values and SEM for all animals. n = 31 mice, 617 sessions.

An important parameter in our task is L_Random_, which dictates the frequency of reversals. Although the animals cannot know the exact value of L_Random_ before each switch because it is drawn randomly, it is possible for the mice to learn the statistical distribution of L_Random_. This knowledge may then be used to infer that the more trials that an animal stays at the better option, the more likely that a block switch might have already occurred. To determine if mice were making use of such knowledge of the task structure, we analyzed the subset of 7,396 blocks (81% of the total of 9,163 blocks) in which animals were adapting quickly after block switches (achieving L_Criterion_ in 20 or fewer trials), therefore focusing on expert-level performance and avoiding periods when animals may be unmotivated. **Figure 1E** shows the histogram of L_Random_ values for these trial blocks, exhibiting the geometric distribution as the task was designed. We found that if L_Random_ was large in the preceding block, the animals tended to choose the better option less frequently prior to the block switch, and subsequently adapted faster after the block switch (**Figure 1F**). The results were qualitatively similar if we included more or all blocks (**Supplementary Figure 1.1**). Moreover, in a smaller number of animals (n = 10 mice, 48 sessions, 312 blocks), we trained them on a variant of the task in which the switching condition is only determined by L_Random_ (i.e., L_Criteiron_ = 0), and the animals exhibited comparable tendency (**Supplementary Figure 1.2**). These results suggest that the mice may anticipate an impending change point and adjust their behavior prior to the block switch.

To quantify the observations, we computed *P (better option) _pre-switch_*, the probability of selecting the better option in the trial immediately before the block switch, and *trials to reach midpoint*, the number of trials from switch for *P (better option)* to reach 0.5. These analyses confirmed the influence of L_Random_ on choice patterns around a block switch (main effect of L_Random_: F (30, 5544) = 6.0743, *P* < 0.001, one-way ANOVA; or if data were binned by 2: main effect of L_Random_: F (15, 5521) = 10.4589, *P* < 0.001; one-way ANOVA; **Figure 1G**) and their speed to adjust after a block switch (main effect of L_Random_: F (30, 711) = 2.1316, *P* < 0.001; or if data were binned by 2: main effect of L_Random_: F (15, 707) = 1.7407, *P* = 0.038; one-way ANOVA; **Figure 1H**). We reiterate that the animal could not predict the exact value of L_Random_ for each block, which was drawn randomly. However, the tendencies around a block switch are consistent with animals learning the task structure, namely the distribution of possible values of L_Random_, presumably through repeated training over dozens of sessions on the two-armed bandit task. As the animal dwells in a block selecting the same better option for many trials, it becomes more probable that a change point in reward probabilities has occurred and therefore the animals should explore the alternate option more. Overall, these results demonstrate that mice were sensitive to the block length – a key feature of the task structure – and could leverage this information during the two-armed bandit task.

### Effects of unilateral lesion of ACAd/MOs on choice behavior around switches

Prior studies implicated the anterior cingulate cortex in behavioral flexibility in the face of variability in the environment (Behrens et al., 2007; Soltani and Izquierdo, 2019). The related region in the mouse is the dorsal aspect of the medial frontal cortex, encompassing the anterior cingulate (ACAd) and medial secondary motor (MOs) areas (Barthas and Kwan, 2017; Laubach et al., 2018; Yang and Kwan, 2021). To determine the role of ACAd/MOs, we trained mice until they reached expert performance, and then performed unilateral excitotoxic lesion by injecting ibotenic acid into the ACAd/MOs region in the left or right hemisphere (n = 5 and 4 mice respectively, 200 pre-lesion and 142 post-lesion sessions in total; **Figure 2A**). For clarity, we will collapse the two groups and refer to trial blocks as ‘lesion’, if the lesioned side was the better option, or ‘contra’ if the side contralateral to the lesion was the better option (**Figure 2B**). Post hoc histology with cresyl violet staining confirmed the loss of cell bodies at the targeted ACAd/MOs location (**Figure 2C – D**). After the lesion, animals performed a similar number of trials and block switches (**Supplementary Figure 2.1**) and had no motor deficit in licking (**Supplementary Figure 2.2)**. Post-lesion mice exhibited block length-dependent choice patterns (**Figure 2E**). However, with the unilateral loss of ACAd/MOs, for the switch from lesion block to contra block, this tendency to choose the worse option pre-switch exacerbated with increasing L_Random_ (left panel, **Figure 2E**). Summary of the data reaffirmed that animals with unilateral ACAd/MOs lesion were selecting the worse option at the expense of exploiting the better option pre-switch, specifically when L_Random_ was large in the preceding block (main effect of lesion: *P* < 0.001, main effect of L_Random_: *P* < 0.001, lesion * L_Random_ interaction: *P* = 0.044, three-way ANOVA; **Figure 2F, Supplementary Table 2.1**). Although they appeared to adapt faster after the switch relative to control animals (main effect of lesion: *P* < 0.001, main effect of side: *P* = 0.003, lesion * side interaction: *P* = 0.038, L_Random_ * side interaction: *P* = 0.004, three-way ANOVA; **Figure 2G**), overall the performance suffered after the lesion (main effect of lesion: *P* = 0.027, main effect of side: *P* = 0.045, main effect of L_Random_: *P* < 0.001, lesion * L_Random_ interaction: *P* = 0.027, three-way ANOVA; **Figure 2H**). The data therefore show that unilateral lesion of the ACAd/MOs impairs the proper estimate and use of task structure knowledge during probabilistic reward learning.

**Figure 2:**
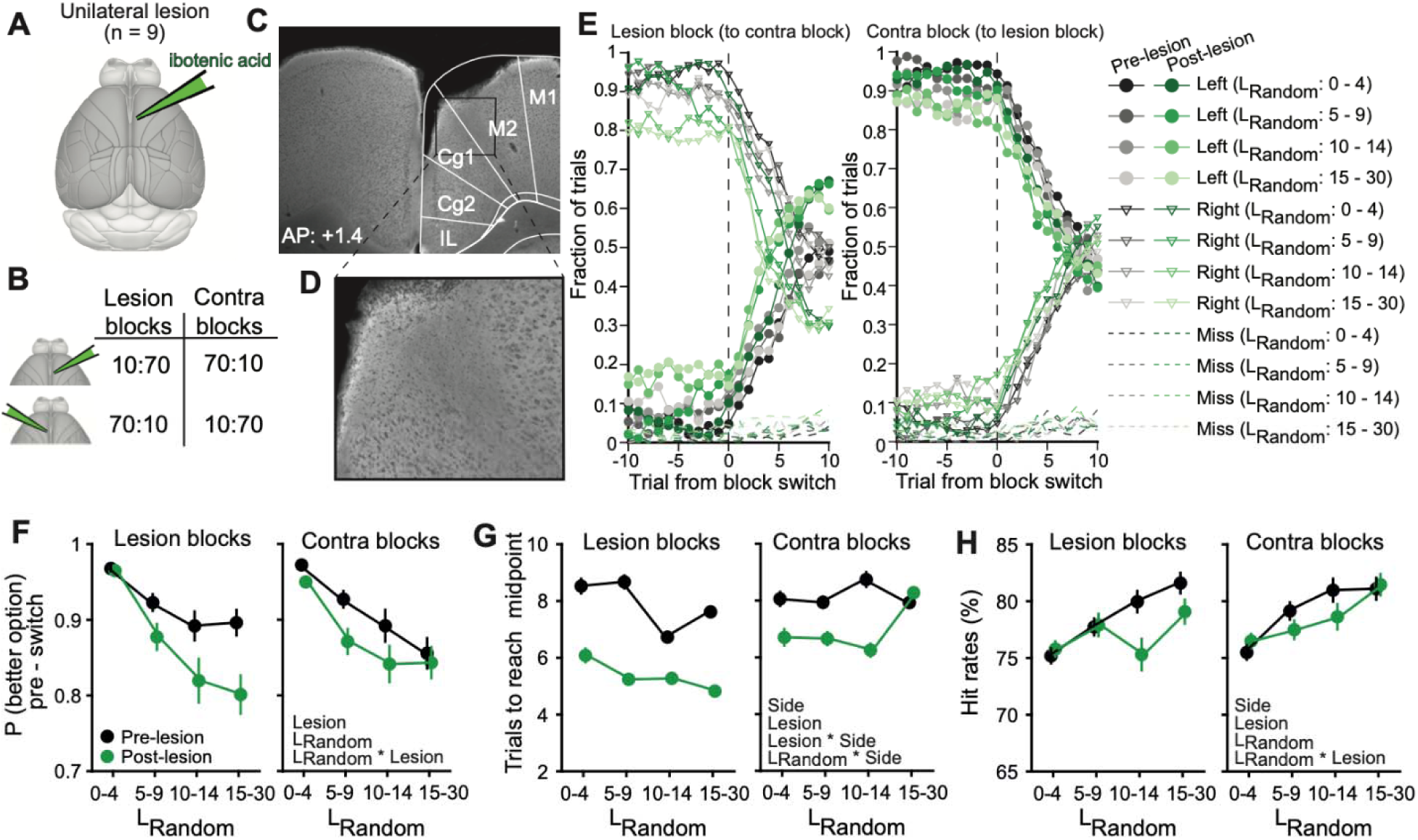
Unilateral lesion of ACAd/MOs altered block-length-dependent choice behavior and impaired overall performance. **(A)** Schematic representation of the unilateral excitotoxic lesion via injection of ibotenic acid. **(B)** Lesion blocks refers to blocks in which the lesioned side is the better option. Contra blocks refer to blocks in which the lesioned side is contralateral to the better option. **(C, D)** Post hoc histology with cresyl violet staining to confirm the loss of neurons in ACAd/MOs. **(E)** Choice behavior around block switches, plotted separately for the 4 ranges of L_Random_. Black, pre-lesion. Green, post-lesion. Left, switches from lesion block to contra block. Right, switches from contra block to lesion block. Mean values and SEM for all animals. **(F)** The probability of choosing the better option on the trial immediately preceding the switch, as a function of L_Random_ for the block preceding the switch. Black, pre-lesion. Green, post-lesion. Mean values and SEM for all animals. **(G)** Similar to (F) for number of trials to reach midpoint (when animal is equally likely to choose either option). **(H)** Similar to (F) for hit rate (probability for animal to choose the better option). For (F) – (H), significant main effects and interactions from three-way ANOVA were indicated (*P* < 0.05). n = 9 mice, 200 pre-lesion sessions and 142 post-lesion sessions.

### A hybrid model of belief and choice kernels to explain the animals’ behavior

To gain insight into the empirical findings, we fitted different computational models to the data. We were specifically drawn to two emerging ideas in the field of decision-making. First, the concept of belief enables an agent to apply their knowledge of the task structure (Jang et al., 2019). Namely, the two-armed bandit task in this study can be conceptualized as a task with two ‘states’ (‘70:10’ and ‘10:70’). For each state, there is an optimal action to take - either choosing left or right, for 70:10 and 10:70 respectively. In this scheme, during each trial, the animal (or agent) holds a belief.

This belief is the probabilities that the task is currently in one of the two states. Then, the agent acts based on this belief. Once the result of the action is revealed, the animal’s belief is updated. This update is influenced by the outcome and the animal’s knowledge of the task structure, which can be approximated as its estimation of the likelihood of a reversal in reward probabilities, which is a change point with hazard rate H. From a Bayesian perspective, the ‘prior’ is the initial belief held by the agent about which state they are in before taking an action. This could be based on their previous experiences or could be a neutral assumption if they have no prior experience. For example, the animal might initially believe it is equally likely to be in either the ‘70:10’ or ‘10:70’ state, or it might have a stronger belief in one state over the other based on past rewards. Second, choice kernels can be used to capture an agent’s tendency to repeat the previous actions (Wilson and Collins, 2019). The choice kernels are updated based on the prior action, scaled by a learning rate. Our belief-CK model contains components for belief and choice kernels, and integrates their outputs for action selection based on a softmax function with separate inverse temperature parameters for belief, and choice kernel, (**Figure 3A, Supplementary Figure 3.1 - 3.4**). Fit to an example session of animal data suggests that this 4-parameter belief-CK model can recapitulate the choice behavior of the mouse in the two-armed bandit task (**Figure 3B – E**).

**Figure 3:**
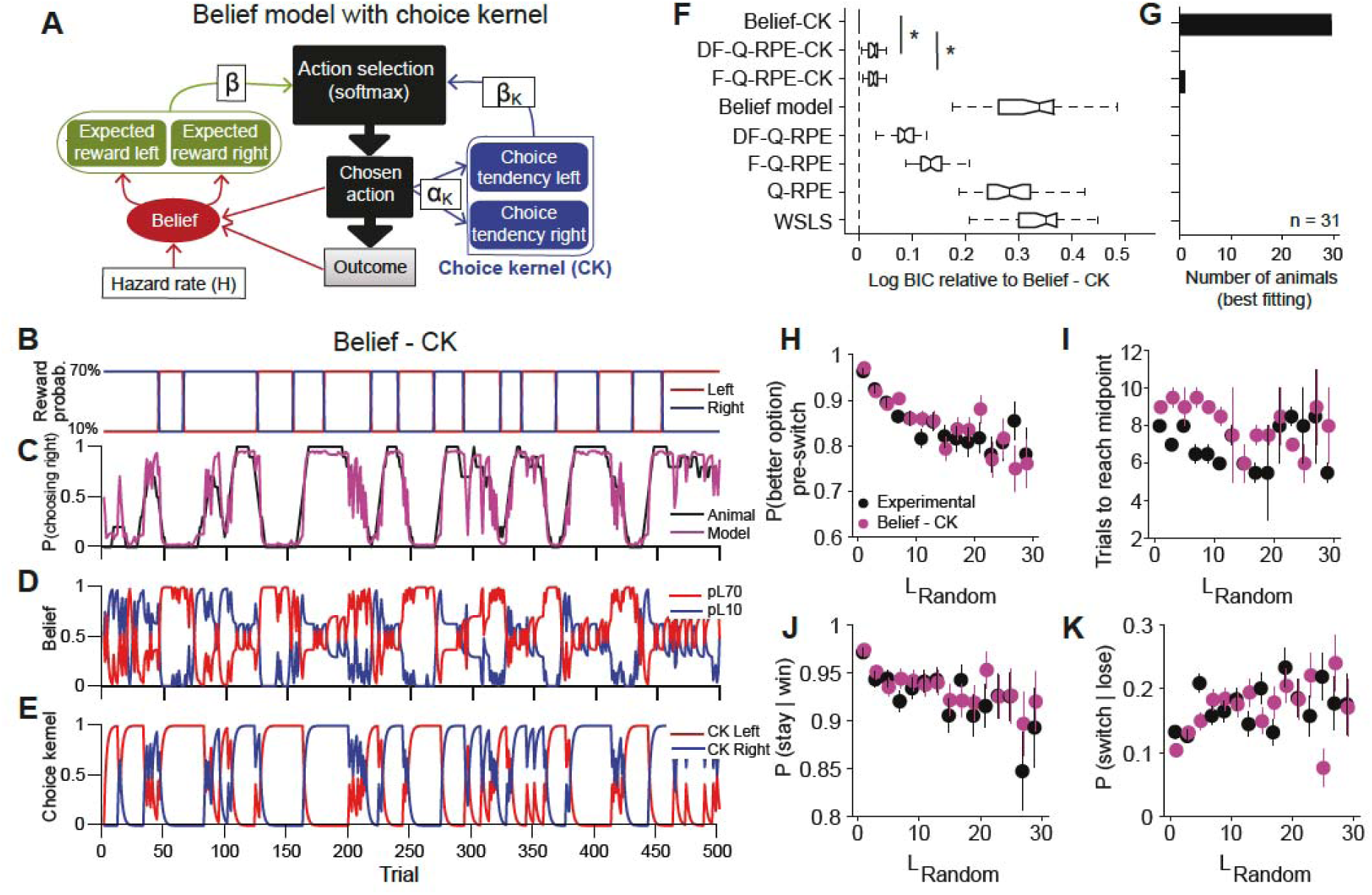
A hybrid model of beliefs and choice kernels to explain the behavior. **(A)** The schematic representation of the belief with choice kernel model (belief-CK). The model has four parameters: H (hazard rate), *β* (inverse temperature for belief), *α_K_* (learning rate for choice kernel) and *β_K_* (inverse temperature for choice kernel). **(B – E)** An example session along with the fits from the belief-CK model, including reward probabilities for left and right options (B) the running-average of probability of choosing right for the animal (black) and model (purple) (C), the belief that the left option is associated with reward probability of 10% (pL10, blue) or 70% (pL70, red) (D), and the choice kernels for left (blue) and right options (red) (E). **(F)** Model comparison between the belief-CK model and 7 other models. Lower log BIC values indicate a better fit. **(G)** The tally of the best-fitting model for each animal. **(H)** The probability of choosing the better option on the trial immediately preceding the switch, as a function of L_Random_ for the block preceding the switch. Black, mice. Purple, simulated performance using the belief-CK model with best-fitting parameters. Mean values and SEM for all animals. **(I)** Similar to (H) for number of trials to reach midpoint (when animal is equally likely to choose either option). **(J)** Similar to (H) for the tendency to win-stay on the 5 trials preceding the switch. **(K)** Similar to (H) for the tendency to lose-switch on the 5 trials preceding the switch. n = 31 mice, 617 sessions.

We compared the belief-CK model against 7 other computational models (see Methods). We started with the win-stay, lose-switch (WSLS) and 3 classic reinforcement learning algorithms including Q-learning (Q-RPE), Q-learning with forgetting (F-Q-RPE), and Q-learning with differential forgetting (DF-Q-RPE) (Ito and Doya, 2015). We then examined effects of adding choice kernels, by testing DF-Q-RPE with choice kernels (DF-Q-RPE-CK), because DF-Q-RPE was the best fit in the initial set of 4 algorithms, and F-Q-RPE with choice kernels (F-Q-RPE-CK), because this model has the same number of free parameters as the belief-CK model. Finally, we also tested the belief model alone without choice kernels. Model comparison based on Bayesian information criterion (BIC) revealed that inclusion of choice kernels improved the fits significantly. Moreover, the belief-CK model had the lowest BIC values (**Figure 3F**; belief-CK versus F-Q-RPE-CK: t_60_= 2.562, *P* = 0.013, paired t-test; belief-CK versus DF-Q-RPE-CK: t_60_= 2.313, *P* = 0.024), and was the best fit for 30 out of 31 animals in this study (**Figure 3G**). For each session, we can simulate the belief-CK model using the best-fitting parameters and compare the tendencies of the simulated and experimental data. This exercise shows that the belief-CK model can capture the L_Random_-dependent choice behavior in the experimental data (**Figure 3H – K**), which is not possible with the classic reinforcement learning algorithm DF-Q-RPE (**Supplementary Figure 3.5**). These analyses demonstrate that simple models of reward-based learning such as Q-learning algorithms cannot fully account for the observed choice behavior. Instead, the results support our intuition that mice were estimating change points, which is formalized as the hazard rate *H* in the belief-CK model.

### Unilateral ACAd/MOs lesion led to side-specific increase in hazard rate for change points

Next, we applied the computational model to quantify the effect of unilateral ACAd/MOs lesion. To account for the possibility of side-specific alterations, we modified the 4-parameter belief-CK model to include 6 parameters to include differential learning for the sides ipsilateral and contralateral to lesion (see Methods; *H*_*lesion*_, *H*_*Contra*_, α_*K lesion*_, α_*K Contra*_, *β*, and *β_K_*). After fitting the expanded model to animal data, we compared pre-versus post-lesion performance in two ways. First, on a per-animal basis, sessions before or after the lesion were concatenated for fitting to yield one set of pre-lesion parameters and one set of post-lesion parameters for each animal. Second, on a per-session basis, each session was analyzed separately and the fitted parameters were summarized.

These analyses revealed a side-specific increase in hazard rate after unilateral ACAd/MOs lesion. The exaggerated hazard rate for the side contralateral to lesion *H*_*Contra*_ was detected on a per-animal basis (**Figure 4A**; pre- vs. post-lesion, *H*_Contra_: *P* = 0.004; post-lesion, *H*_Contra_ vs. *H*_*lesion*_: *P* = 0.810, Wilcoxon signed-rank test), and on a per-session basis (**Figure 4B**; pre- vs. post-lesion, *H_Contra_*: *P* = 0.001; post-lesion, *H_Contra_* vs. *H_lesion_*: *P* < 0.001, Wilcoxon rank sum test). Unilateral ACAd/MOs lesion also led to an increase in choice perseveration for both sides, reflected as higher choice-kernel learning rates (**Figure 4C – D**; per-animal, pre- vs. post-lesion, α_*K lesion*_: *P* = 0.012; α_*K Contra*_: *P* = 0.004; per-session, pre- vs. post-lesion, α_*K lesion*_: *P* < 0.001; α_*K Contra*_: *P* < 0.001, Wilcoxon ranked sum test). Action selection depends on the inverse temperature sum, β+ β_K_, reflecting the exploration-exploitation balance, and inverse temperature ratio, 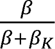 reflecting the relative reliance on belief over choice kernels. There was no detected difference in inverse temperature sum between pre- and post-lesion animals (**Figure 4E – F**, per animal, *P* = 0.567; per session *P* = 0.858, Wilcoxon signed-rank test). By contrast, the inverse temperature ratio was heightened after the lesion (**Figure 4G – H**, per animal *P* = 0.038; per session *P* = 0.001, Wilcoxon signed-rank test). Collectively, these analyses show that the consequences of unilateral ACAd/MOs lesion are a contralateral side-specific increase in hazard rate, and broad increases in choice perseveration and reliance on belief for action selection.

**Figure 4:**
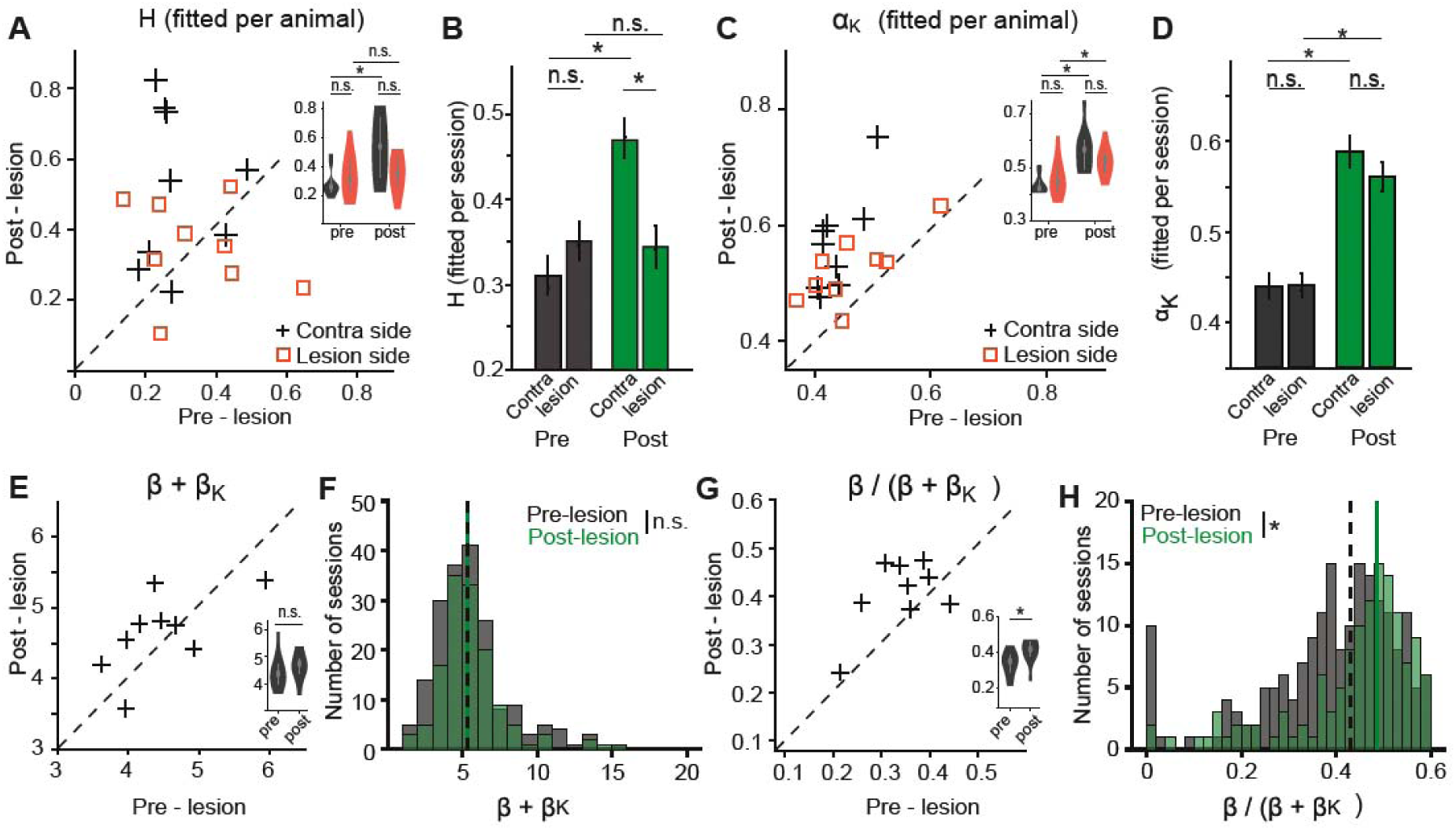
Effects of unilateral lesion of ACAd/MOs is consistent with a side-specific increase in hazard rate. **(A)** The hazard rates, before and after lesion, extracted by fitting the belief-with-choice-kernel model on a per-animal basis. Square, hazard rate for side ipsilateral to lesion. Cross, hazard rate for side contralateral to lesion. Inset, violin plot of the same data. **(B)** The hazard rates, before and after lesion, on a per-session basis. Mean and SEM. **(C – D)** Similar to (A – B) for learning rate for choice kernel. **(E)** The inverse temperature sum, before and after lesion, on a per-animal basis. **(F)** The inverse temperature sum, before and after lesion, on a per-session basis. **(G - H)** Similar to (E – F) for inverse temperature ratio. *, *P* < 0.05. n.s., not significant. n = 9 mice, 190 pre-lesion sessions and 140 post-lesion sessions.

### Accurate change point estimation depends on the balance between the left and right hemispheres

The results so far from unilateral lesions suggest two potential mechanisms for change point estimation. The first possibility is that the computation of change point estimation is lateralized, such that the left hemisphere is involved in estimation for the right side, and vice versa. If this is the case, for a bilateral lesion, we would expect aberrant increases of hazard rates for both sides. The second possibility is that the estimation of change points involves inter-hemispheric coordination, which was perturbed by disruption of one hemisphere. If true, the lack of medial frontal cortex on both sides could nullify their respective maladaptive influences on behavior, and we may observe no or milder deficit after a bilateral lesion of ACAd/MOs. To distinguish between these two possibilities, we injected ibotenic acid bilaterally to the left and right ACAd/MOs regions in expert mice. Animals with bilateral lesions performed fewer trials per session, and accordingly fewer block switches (**Figure 5A – B, Supplementary Figure 5.1;** *P* < 0.001, Wilcoxon signed-rank test), but had no motor deficit (**Supplementary Figure 5.2**). Surprisingly, and in line with inter-hemispheric coordination, there was no detectable change in the L_Random_-dependent choice behavior (**Figure 5C – D**, **Supplementary table 5.1**), and no significant changes in the latent decision parameters including hazard rates (**Figure 5E – H**). Comparison to sham animals in which saline was injected unilaterally (**Figure 5I-P, Supplementary table 5.2**) highlights again that the only effect of bilateral lesion was diminished motivation to perform the task, which was also seen in a prior work from the lab (Siniscalchi et al., 2016). More importantly, these results argue against change point estimation as a lateralized computation in ACAd/MOs, but rather point to unbalance between the hemispheres as the reason for behavioral deficits.

**Figure 5:**
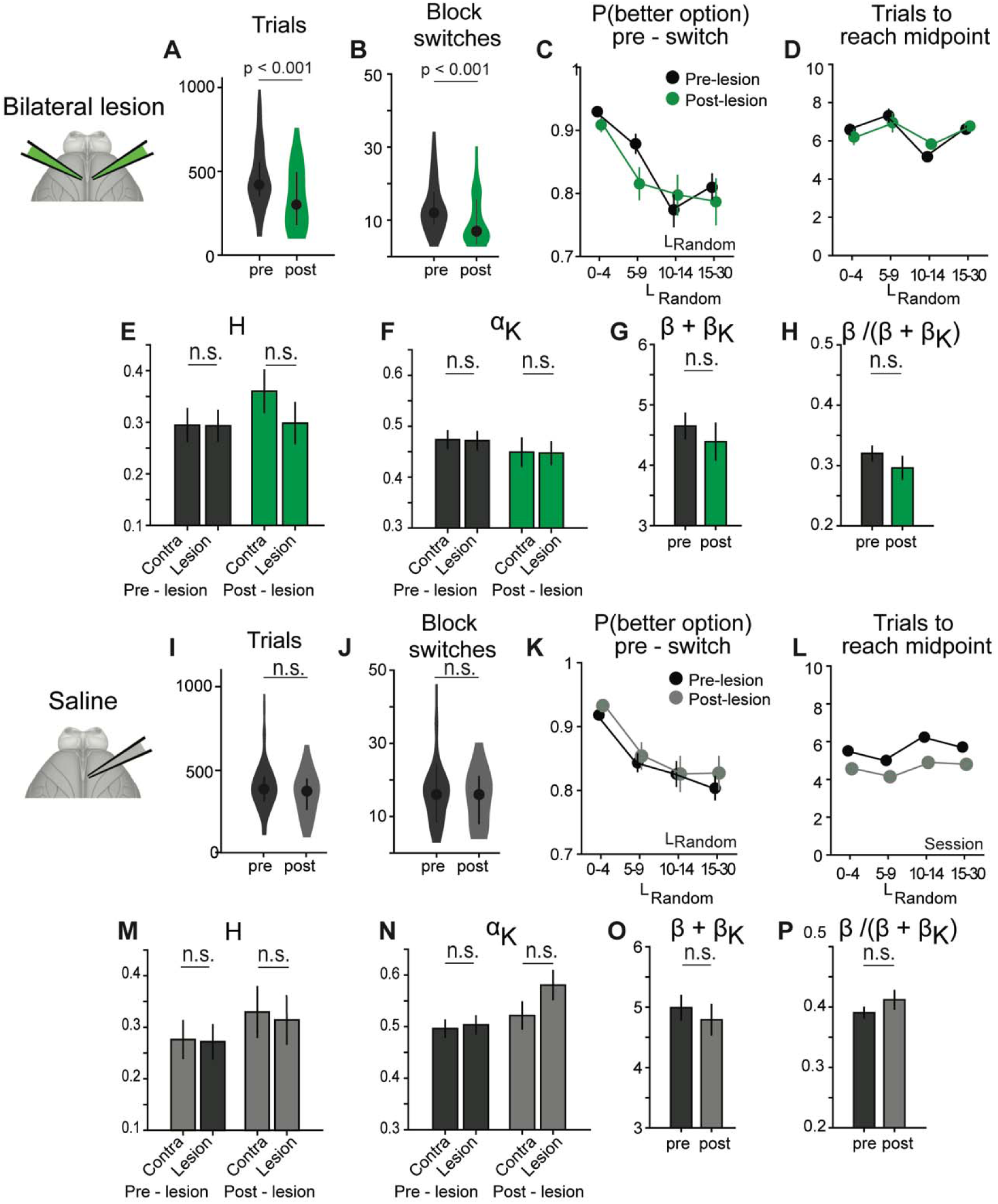
Effects of bilateral and sham lesions of ACAd/Mos. **(A)** The number of trials performed in each session, before and after bilateral lesion, on a per-session basis. Mean and SEM. **(B)** Similar to (A) for the number of block switches in each session. **(C)** The probability of choosing the better option on the trial immediately preceding the switch, as a function of L_Random_ for the block preceding the switch, before and after bilateral lesion, on a per-session basis. Mean and SEM. Significant main effects and interactions from three-way ANOVA were indicated (*P* < 0.05). **(D)** Similar to (C) for number of trials to reach midpoint (when animal is equally likely to choose either option). **(E)** The hazard rates, before and after bilateral lesion, extracted by fitting the belief-CK model on a per-session basis. Mean and SEM. **(F)** Similar to (E) for learning rate for choice kernel. **(G)** Similar to (E) for inverse temperature sum. **(H)** Similar to (E) for inverse temperature ratio. **(I – P)** Similar to (A – H) for sham controls with unilateral saline injection. n.s., not significant. For bilateral lesion, n = 4 mice, 105 pre-lesion sessions and 61 post-lesion sessions. For saline control, n = 4 mice, 117 pre-lesion sessions and 53 post-lesion sessions.

### Medial frontal cortex impacts the decisions during action selection

The lesion-induced effects may be a direct consequence of ACAd/MOs disruption, but some of the behavioral changes can also be due to compensatory adjustments. Therefore, we additionally performed transient inactivation experiments using optogenetics. Mice were implanted with a clear-skull cap that has ∼50% optical transmission (**Supplementary Figure 6.1A**). For photostimulation, we used a laser-steering system (Pinto et al., 2019), in which the excitation beam from a 473 nm laser was steered by a set of mirror galvanometers to specific locations with high spatial and temporal resolutions (**Figure 6A**). We calibrated the linearity of the steered coordinates as a function of galvanometer voltages and the spatial profile of the laser beam (**Supplementary Figure 6.1B – C**). We demonstrated that the system can effectively manipulate neural activity by showing elevated c-fos immunohistostaining after unilateral photostimulation of ACAd/MOs in *CaMKIIa^Cre^;Ai32* animals (**Supplementary Figure 6.1D – F**). Additionally, to determine that our system can effectively bias animal’s behavior, we inactivated the primary visual cortex (V1) and anterolateral motor cortex (ALM) by photostimulating parvalbumin-expressing interneurons in *Pvalb^Cre^;Ai32* animals. No effect was observed when silencing V1, whereas biased tongue licks were induced by inhibiting ALM in mice during the two-armed bandit task, consistent with previous findings (Guo et al., 2014) **(Supplementary Figure 6.2).**

**Figure 6:**
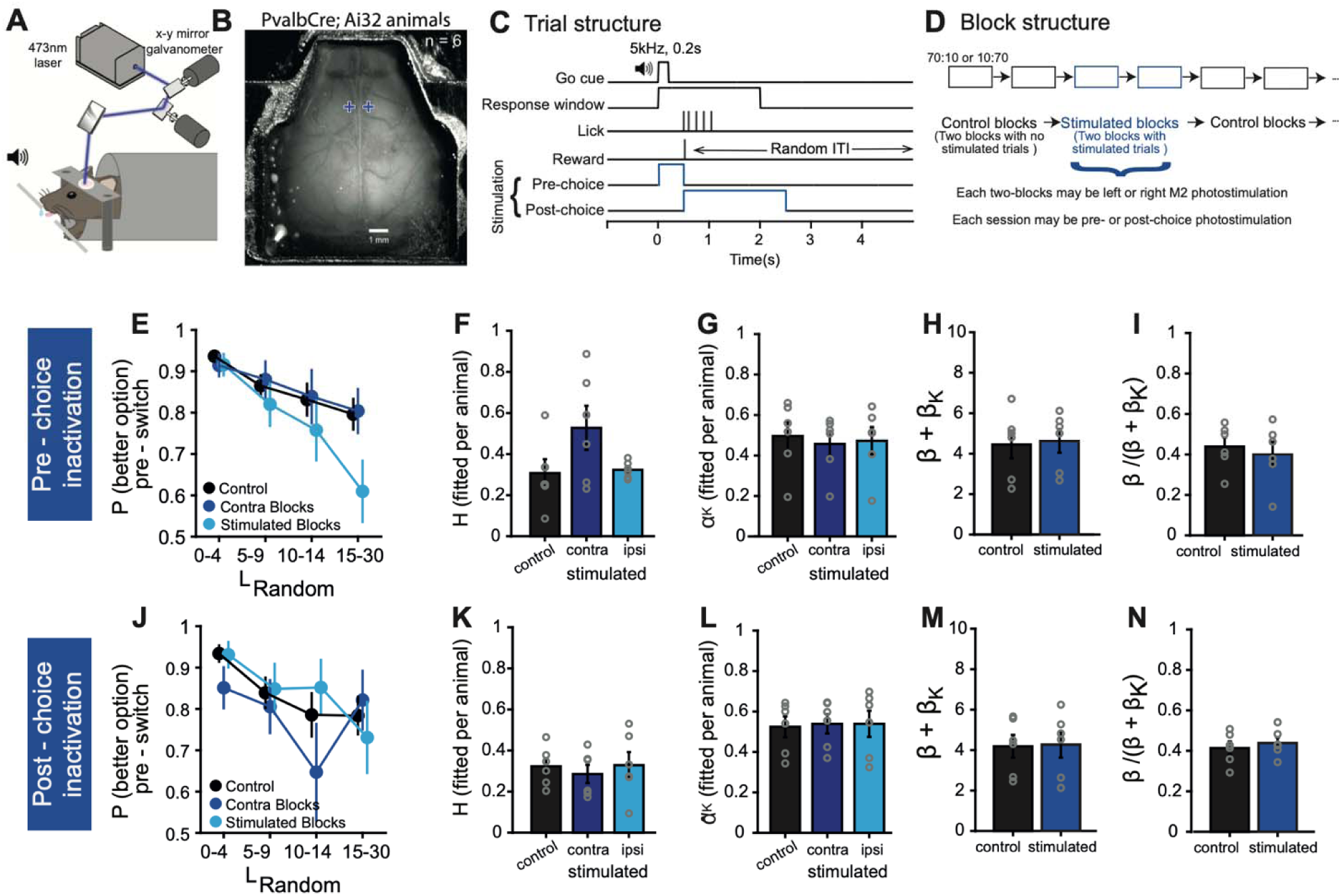
Optogenetic inactivation in pre-choice, but not post-choice, period reproduced the deficit in change-point estimation. **(A)** The schematic representation of experimental setup. **(B)** CCD image of a mouse with a cleared skull cap. The tw blue crosses indicate the locations of the photostimulation, i.e. left and right ACAd/MOs. **(C - D)** The trial and block structures, and the timing of the photostimulation. **(E)** The probability of choosing the better option on the trial immediately preceding the switch, as a function of L_Random_ for the block preceding the switch for pre-choice inactivation. Black, control blocks. Light blue, Stimulated blocks. Blue, contralateral to stimulated blocks. Mean values and SEM for all animals. **(F)** The hazard rates extracted by fitting a modified belief-CK model, for pre-choice inactivation, on a per-animal basis. **(G)** Similar to (F) for learning rate for choice kernel. **(H)** Similar to (F) for inverse temperature sum. **(I)** Similar to (F) for inverse temperature ratio (**J-N**) Similar to E-I for post-choice inactivation. n = 6 animals.

To suppress excitatory activity in ACAd/MOs during block switches, we used *Pvalb^Cre^;Ai32* animals in which the channelrhodopsin ChR2 was selectively expressed in parvalbumin-expressing (PV) GABAergic interneurons (n = 6). Targeted photostimulation would activate PV interneurons in the left or right ACAd/MOs (**Figure 6B**), which would in turn silence local excitatory spiking activity (Guo et al. 2014; Li et al. 2019). These transient inactivations were applied either at the time of action selection (“pre-choice”, from cue onset to lick response) or after the outcome (“post-choice”, from lick response for 2 s; **Figure 6C**), and on every trials across two consecutive blocks such that activity was suppressed before and after certain block switches (**Figure 6D**).

Pre-choice inactivation impaired the ability of the mice to select the better option in the trials immediately preceding the block switch, when the stimulated side was contralateral to the side with the better option (**Figure 6E**), suggesting a lateralized influence of ACAd/MOs on decision-making under conditions of uncertainty as observed in lesion data. In contrast, post-choice inactivation did not produce significant changes in the selection of the better option immediately before block switches (**Figure 6J**, indicating that the effects of ACAd/MOs inactivation are primarily restricted to the pre-choice period (three-way ANOVA: main effect of block length: F(3, 1999) = 49.778, *P* < 0.001; main effect of stimulation period: F (1, 1999) = 7.974, *P* = 0.004; interaction between block length and stimulation: F(3, 1999) = 2.674, *P* = 0.046; interaction between block length, stimulation and stimulation period: F(3, 1999) = 2.625, *P* = 0.049).

Fitting to the belief-CK model, expanded to account for the optogenetic stimulation (see Methods; *H_Control_, H_ipsi stim_, H_Contra stim_, α_K Control_, α_K ipsi stim_, α_K Contra stim_, β_Control_, β_stim_, β_K Control_* and *β_K stim_*), highlights the strongest effect is an acute change to hazard rate contralateral to the transient inactivation for pre-choice inactivation (**Figure 6F**; pre-choice inactivation, *H_Contra stim_* vs. *H_ipsi stim_*: *P* = 0.156; *H_Contra stim_* vs. *H_Control_*: *P* = 0.094; **Figure 6K**; post-choice inactivation, *H_Contra stim_* vs. *H_ipsi stim_*: *P* = 0.687; *H_Contra stim_* vs. *H_Control_*: *P* = 0.562, Wilcoxon signed-rank test), although there were variations across individual animals and effect was not statistically significant There were no detectable effects of transient inactivation on 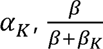, and *β + β_k_* for pre-choice inactivation (Figure 6G-I) or post-choice inactivation (**Figure 6K-N**). Together with the results from unilateral lesions, we interpret these findings from acute inactivation to indicate that ACAd/MOs is involved specifically in the change point estimation process, which occurs during the pre-choice period.

## Discussion

This study provides evidence that mice anticipate impending change points by altering their choices prior to switches in a classic probabilistic reward learning task. Computational analyses indicate that the animals’ choice behavior is consistent with a model of belief updating and choice perseveration. Causal perturbation experiments emphasize the role of the ACAd/MOs region of the medial frontal cortex. Crucially, as discussed below, the collective results from the range of manipulations employed – unilateral and bilateral lesions, as well as pre- and post-choice optogenetic inactivation – provide important insights that can constrain the potential neural mechanisms underlying change-point estimation during decision-making.

Although many studies of decision-making in rodents relied on analyses involving Q-learning algorithms, there are other reports suggesting deviations from simple reinforcement learning. For instance, a pioneering study demonstrated that rats are exceedingly sensitive to changes in reward rates, approximating an ideal observer (Gallistel et al., 2001). Moreover, several studies using a variety of timing, operant conditioning, decision, and sensory categorization tasks have found neural and behavioral data consistent with the use of belief in rodents (Karlsson et al., 2012; Li and Dudman, 2013; Liu et al., 2021; Starkweather et al., 2017; Vertechi et al., 2020). However, these prior studies are different because in some cases, the use of belief did not necessarily confer better performance over other strategies for the task (Starkweather et al., 2017). In other cases, the task employs a series of deterministic outcomes (Vertechi et al., 2020), which strongly favors switching behaviors. Our study therefore extends these past results by showing decisions consistent with belief updating in one of the most popular value-based decision-making tasks used for human and animal studies (Sutton and Barto, 2018).

To quantify the animals’ behavior, we proposed a model involving belief updating with a fixed hazard rate, which was motivated by a prior study (Jang et al., 2019) and adapted to fit our task design. In this model, the agent understands that each action is associated with one of two reward probabilities. Given information from its choice and reward history as well as knowledge of the probability of a switch in reward probabilities, the agent infers the likelihood of the current states associated with the actions. This is in sharp contrast to the Q-learning algorithms, where the agent is implicitly ignorant of the task structure, and simply updates action values based on the last trial’s action and outcome. For belief updating, one limitation for our model is that the hazard rate is a constant value within a session. This assumption seems reasonable because mice were trained on the task extensively and probably accrue knowledge of the hazard rate based on experience of multiple switches across many sessions. That said, in principle, it is possible for an agent to infer and adjust the hazard rate as the task proceeds (Wilson et al., 2010). Moreover, there are other models, such as ones based on a flexible learning rate, that can account for variable choice behavior as a function of outcome history (Grossman et al., 2022). Future studies may employ tasks that can more specifically disambiguate between the different ways in which mice may adapt learning parameters within a session.

To determine the role of the medial frontal cortex, we used both permanent lesions and transient inactivation, methods that each have its advantages and limitations (Otchy et al., 2015; Vaidya et al., 2019) and they provide complementary insights into the functioning of this brain region. Specifically for lesions, we employed unilateral manipulations, such that we can compare effects between sides ipsilateral and contralateral to the lesion in the same animal, serving as a rigorous internal control. Results from these experiments demonstrated lateralized deficits from lesions of the medial frontal cortex. This finding may be surprising, because although sensory and motor functions are typically expected to be lateralized, it is less obvious that cognitive function may also be side-dependent. However, we note that a few prior studies have also found side-specific decision-making deficits from unilateral manipulations, such as effects of dorsal striatum on action values (Tai et al., 2012) and effects of frontal cortex on lapses in a multisensory task (Pisupati et al., 2021) and short-term memory (Yin et al., 2022). There were also instances of hemi-neglect in humans (Stone et al., 1991; Kerkhoff, 2001; Crowne et al., 1986; Reep and Corwin, 2009). The exact reason for side-specific effects is unclear, although one possibility is that when decisions are intimately tied to responses associated with lateralized motor actions, then there is embodiment and motor and premotor cortical regions become involved in the neural computation (Bennur and Gold, 2011).

The various deficits arising from lesion and optogenetic manipulations are useful for thinking about the potential mechanisms for how the medial frontal cortex contributes to belief updating and more specifically change-point estimation. An inaccurate estimate, which would reflect as altered hazard rate in our computational model, can occur for several reasons: (1) error in estimating the value of hazard rate, (2) error when using the hazard rate to update belief, and (3) error when using the prior choice and reward to update belief. Reason (3) was not explicitly tested in our model fits but could manifest as an apparent change in hazard rate. Among these possibilities, the first option seems unlikely. In our task, an accurate value for hazard rate cannot be determined quickly but must be calibrated by experiencing many switches across multiple sessions. This is difficult to reconcile with the immediate deficit observed with pre-choice optogenetic inactivation. The third option is also unlikely. Previous studies have shown that choice- and outcome-related signals arise in the medial frontal cortex shortly after the outcome (Siniscalchi et al., 2019; Sul et al., 2011), whereas optogenetic inactivation during this post-choice period was ineffective. Therefore, it may be the case that the medial frontal cortex is involved in incorporating the likelihood of an impending change point for estimating the current task state.

Furthermore, rather than computing using a probability such as the hazard rate, the animal may instead approximate the process by employing simpler heuristics to predict the impending occurrence of a change point. One intuitive heuristic, consistent with the reason for lateralized deficits, is that the animals may rely on the recent choice history of the number of better options chosen, which would indicate a higher likelihood of an impending switch. Here, the lack of effect from bilateral lesions can shed light on the form of the heuristic. For example, one heuristic that can work is a ratio of the number of recent left choices divided by the number of recent right choices, and if the unilateral lesion effectively adds a multiplier to the side’s choice history, then a bilateral perturbation would lead to a null effect. Heuristics based on choice history are plausible because the medial frontal cortex has long-lasting, persistent representation of past choices (Bari et al., 2019; Hattori et al., 2019; Siniscalchi et al., 2016; Sul et al., 2011). One caveat for this line of logic is that it is based on a specific belief updating model. However, we discuss the implications to illustrate how the results can inform the underlying neural basis.

To sum, the two-armed bandit task has gained widespread use in neuroscience and artificial intelligence research because of its simplicity, translational significance, and amenability to computational modeling. Our results show that mice may perform the task by not only updating based on choices and outcomes, but also leverage knowledge of the environment to estimate change points. The diminished or exaggerated use of this prior knowledge represents suboptimal decision-making, which may underlie pathological behaviors in neuropsychiatric disorders that involve dysfunctions of the medial frontal cortex.

## METHODS

### Lead Contact

Further information and requests for resources and reagents should be directed to and will be fulfilled by the Lead Contact, Huriye Atilgan

### Materials Availability

All published reagents and mouse lines will be shared upon request within the limits of the respective material transfer agreements. Detailed plans including parts list for constructing the behavioral training apparatus is available at https://github.com/Kwan-Lab/behavioral-rigs.

### Data and Code Availability

Data and analysis software for this paper will be available at Github (https://github.com/Kwan-Lab).

## EXPERIMENTAL MODEL AND SUBJECT DETAILS

### Mouse lines

In this study, we used a total of 30 adult male mice (**Table 1**; 2 - 8 months old), including 24 C57BL/6J wild-type mice (#000664, Jackson Laboratory) for the lesion experiments, and 6 *Pvalb^Cre^*;*ROSA^CAG-ChR2-EYFP^(Ai32)* mice for the photostimulation experiments. The *Pvalb^Cre^*;*ROSA^CAG-ChR2-EYFP^(Ai32)* mice were generated by crossing the *Pvalb^Cre^*(B6.129P2-*Pvalb*^tm1(cre)Arbr^/J; #017320, Jackson laboratory) and *ROSA^CAG-ChR2-EYFP^(Ai32)* (B6.Cg-*Gt(ROSA)26Sor^tm32^ ^(CAG-COP4*H134R/EYFP)Hze^*/J; #024109, Jackson laboratory) strains. Mice were housed in groups of 2 – 5 per cage in a 12h:12h light:dark cycle with ad libitum access to food. All of the experiments were completed during the light cycle. Experimental procedures were approved by the Yale University Institutional Animal Care and Use Committee.

## METHOD DETAILS

### Surgery for lesion experiments

All of the mice in the lesion study underwent two surgeries. In the first surgery, a stainless steel headplate was attached to the skull to facilitate behavioral training. After collecting baseline behavioral data, a second surgery consisting of either an excitotoxic or sham lesion was performed. Before each surgery, the animal was treated pre-operatively with carprofen (5 mg/kg, i.p.; 024751, Butler Animal Health) and dexamethasone (3 mg/kg, i.p.; Dexaject SP, #002459, Henry Schein Animal Health). At the start of each surgery, anesthesia was induced with 2% isoflurane in oxygen, and the animal was placed on a water-circulating heating pad (TP-700, Gaymar Stryker). The head was secured in a stereotaxic frame with ear bars (David Kopf Instruments). Following induction, isoflurane concentration was lowered to 1 – 1.5% based on the animal’s weight and breathing pattern.

For the first surgery, the scalp was shaved using scissors and cleaned with povidone-iodine (Betadine, Perdue Products L.P.). A narrow portion of the scalp was removed along the midline from the interaural line to a line visualized just posterior to the eyes. The scalp was retracted to expose the dorsal aspect of the skull and washed thoroughly with artificial cerebrospinal fluid (ACSF; in mM: 5 KCl, 5 HEPES, 135 NaCl, 1 MgCl2, and 1.8 CaCl2; pH 7.3). A scalpel and a ballpoint pen were used to scratch and paint marks onto the skull at the secondary motor and anterior cingulate cortices (MOs/ACAd; +1.5 mm AP, +0.3 mm ML from bregma), to be used as a landmark for the second surgery. A custom-made stainless-steel head plate (eMachineShop) was then bonded to the skull with cyanoacrylate glue (Loctite 454, Henkel) and transparent dental acrylic (C&B Metabond, Parkell Inc.), with care taken to cover any remaining exposed skull. The post-operative care was provided immediately, and for three consecutive days following surgery, consisting of carprofen (5 mg/kg, i.p.) for analgesia and preservative-free 0.9% NaCl (0.5 mL, i.p.) for fluid support. The animal had at least one week for post-operative recovery prior to the onset of behavioral training.

For the second surgery, a 1-mm-diameter circular craniotomy was made over the marked spot using a high-speed rotary drill (K.1070, Foredom). A total of ∼300 nL of ibotenic acid (5 mg/mL in saline; 505024, Abcam) was injected into two locations (+1.5 mm and +1.7mm AP, +0.3 mm ML from bregma; 0.4 mm DV) through a glass micropipette attached to a microinjection unit (Nanoject II, Drummond). More specifically, each location would receive 15 pulses of 9.6 nL of the prepared solution. To minimize backflow of the injected solution, there was a 1 min gap between each pulse, and the micropipette was left in place for 20 min after the last pulse. Sham animals underwent the same surgical procedure, but saline was delivered instead of ibotenic acid. The exposed skull was covered with dental cement. The animal had two weeks of post-operative recovery prior to resuming behavioral testing.

### Surgery for photostimulation experiments

All of the mice in the photostimulation study underwent one surgery. The animal was anesthetized in the same way as described above. Procedures to prepare the skull were nearly identical to those described in (Pinto et al., 2019). Briefly, the scalp covering the dorsal skull surface was excised and the periosteum over the skull was removed using a micro-curette (VWR Buck Micro Curette, 10806-346). The skull was washed thoroughly with ACSF. A custom stainless-steel head-plate (eMachineShop) was affixed at points above the cerebellum and olfactory bulbs with cyanoacrylate glue (Loctite 454, Henkel) and transparent dental acrylic (C&B Metabond, Parkell Inc.). The exposed skull was covered with a thin layer of cyanoacrylate glue (Apollo 2000, Cyberbond) and transparent dental acrylic, then polished with an acrylic polishing kit (0321, PearsonDental), and finally covered with transparent nail polish (72180, Electron Microscopy Services). The animal had at least one week of post-operative recovery prior to the onset of behavioral training.

### Behavioral training apparatus

The apparatus for training head-fixed mice was adapted from (Siniscalchi et al., 2016). Detailed plans including parts list for constructing the behavioral training apparatus is available at https://github.com/Kwan-Lab/behavioral-rigs. Briefly, the behavioral box was constructed using a closed compartment of an audio-visual cart (4731T74, McMaster-Carr) that was soundproofed with acoustic foam (5692T49, McMaster-Carr). The mouse was placed in an acrylic tube (8486K433, McMaster-Carr), which allowed for postural adjustments but restricted large body movements. Two metal screws were used to attach the head plate of the mouse onto a custom stainless-steel mount (eMachineShop). The lickometer was based on a 3D-printed part that held two lick ports constructed from 20-gauge needles, and was placed in front of the mouse such that the lick ports are on the left and right of the animal’s mouth. The position of the lick ports relative to the mouse could induce considerable side bias and influence response time. To mitigate variations across sessions, the lickometer was attached to an XYZ translation stage (MT3, Thorlabs) for precise positioning, and the same set of coordinates were used for the same mouse between sessions.

Water was supplied to the lick ports via Tygon tubing (EW-95666-01, Cole-Parmer). A touch detector circuit was used for detecting tongue licks onto each lick port. Water was delivered at the lick ports by gravity feed and controlled by solenoid valves (EV-2-24; Clippard or MB202-V-A-3-0-L-204, Gems Sensor Solenoid). The water amount is controlled by the duration of a TTL pulse, and we calibrated the solenoid to deliver ∼2 *μ*L per pulse. All of the electrical circuits for water delivery and lick detection were connected to a desktop computer via a data acquisition board (USB-201, Measurement Computing). A pair of speakers (S120, Logitech) were positioned in front of the animal for auditory stimuli (calibrated to 80 dB). The tasks were programmed in scripts using the Presentation software (Neurobehavioral Systems), which controlled the entire behavioral apparatus including stimulus presentation. A table lamp (LT-T6, Aukey) was placed in each box, behind the mouse, to provide dim ambient light in the box. A camera (SV-USBFHD01M-BFV, Svpro) was used to optimize the lick port position at the beginning of each session and monitor the animal’s behavior throughout the session.

### Two-armed bandit task

Mice were fluid-restricted. On training days, animals received all of their water intake from behavioral training that occurred 1 session per day, 5 days per week. On non-training days and days when weight measurements fell below 85% of their pre-restriction weight, water was provided ad libitum in the home cage for 5 minutes.

Prior to any behavioral training, the animal was handled and habituated to head fixation for increasing durations over three days. Water was manually provided via the lick ports to familiarize mice with receiving fluid from the lickometer. After 1 – 2 days of habituation, the animal underwent two phases of shaping.

In the first phase, the animal was trained to alternate between the two lick ports to receive water rewards. More specifically, on each trial, there would be an auditory cue (duration = 0.2 s, tone with 5 kHz carrier frequency). The onset of the auditory cue is the start of a 5-s long response window, during which the first lick detected is the animal’s response. The playback of the auditory cue was terminated early if the response was recorded before the entire stimulus was played. The animal was required to alternate between left and right responses to earn water rewards: if the last rewarded response was left, then the mouse must make a right response to receive water, and vice versa. The inter-trial interval had a fixed duration, such that the auditory cue for the next trial would occur 3.1 s after the animal’s response. A session would end when the animal did not lick during the response window (‘miss’) for 20 consecutive trials. When the animal could attain at least ∼60 rewards in a session, the shaping would proceed to the second phase.

In the second phase, the animal still had to alternate, but was trained to the trial structure including withholding licks between trials. The second phase was similar to the first phase, with two exceptions. First, the onset of the auditory cue is the start of a 2-s long response window, during which the first lick detected is the animal’s response. Second, the addition of a no-lick period between trials. The no-lick period began 3 s after the animal’s response. Initially, the duration of the no-lick period was drawn from a truncated exponential distribution (λ = 0.33333, minimum = 1, maximum = 5). If any lick was detected during the no-lick period, then another duration drawn from the same truncated exponential distribution would be added onto the end of the first no-lick period. The addition could repeat for up to 5 times if the animal could not withhold licking. Therefore, the possible duration for the entire no-lick period ranged between 1 and 25 s, and was dependent on whether the animal could withhold licking. Subsequently, the auditory cue for the next trial would occur 0.1 s after the end of the no-lick period. When the animal could receive rewards in at least ∼40% of all trials, it would be advanced to the two-armed bandit task. Typically, the animal would proceed through each shaping phase in 3 or fewer sessions.

In the two-armed bandit task, the auditory stimulus, response timing, and inter-trial interval including no-lick period were exactly the same as the second shaping phase. However, the outcome of each trial was probabilistically determined. In a 10:70 block of trials, the left lick port had a 10% chance of delivering water if chosen and the right lick port had a 70% chance of delivering water if chosen. By contrast, in a 70:10 block of trials, the reward probabilities associated with the left and right ports were reversed. Hence, the better option is right in a 10:70 block, but left in a 70:10 block. At the start of each session, the block type (10:70 or 70:10) was randomly chosen.

The block type would switch when the mouse fulfilled the switching condition: perform trials (L_Criterion_) until the animal accumulated 10 choices selecting the side with high reward probability, and then perform additional trials (L_Random_) that were drawn from a truncated geometric distribution (*p* = 0.0909, no minimum = 0, maximum = 30). Notably, L_Criterion_ depended on the animal’s performance, whereas L_Random_ was random and independent of performance. The block type would continue to switch, as long as the animal was fulfilling the switching condition of each block. In the lesion experiments, they would be tested on the two-armed bandit task in daily sessions until at least 150 switches were collected for each of the pre- and post-lesion conditions.

### Photostimulation

The photostimulation rig allowed for rapid adjustment of the position of the laser. The rig was constructed based on the design in (Pinto et al., 2019). Briefly, a 473 nm laser beam (Obis LX 473 nm, 75 mW; 1193830, Coherent) was steered by a set of XY galvo mirrors (6210H, Cambridge Technologies) mounted in a ThorLabs 60 mm cage system. The laser was sent through a F-theta scan lens (f = 160 mm; FTH160-1064-M39, ThorLabs) and directed onto the animal’s head. A monochromatic camera (Grasshopper3; GS3-U3-23S6M-C, Point Grey) equipped with a telecentric lens (TEC-55, Computar) was used to visualize the cortical surface and to calibrate the position of the laser beam relative to bregma. The laser, mirrors, and camera were controlled via a data acquisition board (PCIe-6343, National Instruments) by custom software written in MATLAB on a desktop computer. The laser was calibrated to yield a time-averaged power of 1.5 mW at the sample. Light transmission through the clear-skull cap (dental cement and skull) was measured by placing the cap at the sample plane, and positioning a laser power meter underneath the cap.

Animals underwent the same shaping phases and task training. For the photostimulation experiments, the animal was tested on the two-armed bandit task in a behavioral setup within the photostimulation rig. Temporally, the photostimulation could occur either before or after the animal’s response. For pre-choice photostimulation, the laser was turned on at the onset of the auditory cue and turned off immediately when a response was detected. For post-choice photostimulation, the laser was turned on immediately when a response was detected and turned off 2 s later. Spatially, the photostimulation was targeted to one of two possible locations: left MOs/ACAd (+1.5 mm AP, - 0.3 mm ML from bregma) or right MOs/ACAd (+1.5 mm AP, +0.3 mm ML from bregma).

At the start of each session, the timing of the photostimulation (pre-choice or post-choice) was randomly chosen and stayed the same for the entire session. The initial 3 – 5 blocks were always control blocks, i.e., no photostimulation. The rationale was to make sure the animal was performing the task well that day before any perturbation. Subsequently, the next 2 blocks would be photostimulation blocks targeting the same spatial location, followed by 2 control blocks, followed by 2 photostimulation blocks targeting the same spatial location, and so on. For the photostimulation blocks, the spatial location was randomly selected to be left or right MOs/ACAd each time. In other words, in the same session, the animal may receive perturbation of both left and right MOs/ACAd, albeit in different trial blocks.

To prevent the animal from using stray laser light to distinguish photostimulation from control blocks, we implemented a masking stimulus by shining a blue LED at the eyes. The masking stimulus had the same onset timing and duration as the photostimulation used for the session, and was applied for every trial in both control and photostimulation blocks.

### Histology

To determine the extent of the lesions, following behavioral experiments, the mouse was deeply anaesthetized with an overdose of isoflurane and transcardially perfused with chilled formaldehyde solution (4%, in phosphate-buffered saline (PBS)) at a rate of 5 mL/min. The brain was quickly removed, stored overnight in the formaldehyde solution at 4 °C, and then switched to PBS for long-term storage. Coronal sections with a thickness of 100 μm were cut using a vibratome (VT1000 S, Leica).

For cresyl violet staining, cresyl violet (1 g/L; 10510-54-0, Sigma Aldrich) was added to filtered *H*_2_O and stirred overnight. The next day, glacial acetic acid (2.5 mL/L; 64-19-7, Sigma Aldrich) was added to the solution. The tissue sections were washed with filtered *H*_2_O before mounting on glass slides and stained with a pre-warmed (50°C) cresyl violet solution. The sections were dehydrated with ascending grades of alcohol (95% for 10 minutes, 100% twice for 10 minutes each), cleared with xylene (twice for 5 minutes each), and mounted with DPX mounting medium (06522, MilliporeSigma).

For NeuN staining, tissue sections were washed three times with PBS and then incubated with a blocking solution (5% normal goat serum, 0.3% Triton X-100, in PBS) for 1 hour at room temperature. Subsequently, sections were incubated with rabbit monoclonal primary antibody against NeuN (1:500 dilution; ab177487, Abcam Inc,) overnight at 4□°C on the shaker. After washing three times with PBS, tissue sections were incubated with goat anti-rabbit secondary antibody with conjugated Alexa 488 (1:500 dilution; ab150077, Abcam Inc,) for 2.5 hours at room temperature. After washing with PBS, nuclear staining was performed by incubating with a 4′,6-diamidino-2-phenylindole (DAPI) staining solution (ab228549, Abcam Inc.) for 10 minutes. Finally, sections were washed three times with PBS and then with filtered *H*_2_O, before mounting on slides with DPX mounting medium (06522, MilliporeSigma). A motorized upright fluorescence microscope (Olympus BX61, Olympus) was used to image the sections.

## QUANTIFICATION AND STATISTICAL ANALYSIS

### Analysis of behavioral data

Timestamps of the behavioral events, including cue onsets, outcome onsets, licks, and reward probabilities were logged to a text file by the NBS Presentation software. The text files were parsed and analyzed using scripts written in MATLAB (MathWorks, Inc.). For all of the analyses, we excluded the session if the animal had fewer than 4 block switches. We analyzed all of the trials up to the last switch, and ignored the trials in the last incomplete block where by definition had many miss trials.

When analyzing the consequences of unilateral lesions, for simplicity, we used the term *<u>lesion</u> <u>blocks</u>* and *<u>contra blocks</u>*. This is because unilateral lesions were randomly assigned to the left or right hemisphere for each animal. Lesion blocks refer to those blocks where the lesioned side is the same as the better option. In other words, if the animal had a unilateral lesion on the right hemisphere, then the lesion blocks correspond to 10:70 blocks. If the animal had a unilateral lesion on the left hemisphere, then the lesion blocks correspond to the 70:10 blocks. The remainder was referred to as the contra blocks.

### Analysis of behavioral data – effects of block length

For analyses involving block lengths, we used the subset of data in which L_Criterion_≤ 20 trials for the pre-switch block, in order to restrict the analyses to situations where the performance was similar.

The probability of choosing the better option pre-switch*, <u>P</u>* <u>(*better option*) _pre-switch_</u>, was determined for each animal by examining the last trial before each block switch, dividing the number of times in which the animal chose the initial better option (i.e., the side with 70% reward probability before switch) by the number of switches. <u>Hit rate</u> was the proportion of trials in which the animal selected the better option. The win-stay probability, <u>P (stay | win)</u>, was the fraction of trials in which animals repeated a choice after a rewarded trial. The lose-switch probability, <u>P (switch | lose)</u>, was the fraction of trials in which an animal switched its choice after an unrewarded trial. For all of these performance metrics, we computed the metric on a session-by-session basis, then averaged across sessions to obtain per-animal value. In the lesion data, <u>P (better option) _pre-switch_</u>, <u>P (stay |</u> <u>win)</u> and <u>P (switch | lose)</u> was calculated in the last five trials before each block switch

<u>Trials to reach midpoint</u> was determined by first calculating the fraction of trials for choosing the initial better option around the block switch for the animal, and then identifying the trial from the switch where the fraction of trials choosing the initial better option was closest to 0.5 To compute the trials to reach the midpoint metric, we would first concatenate data across sessions including inserting 20 NaN in the gaps, then compute the metric to obtain the per-animal value. This allowed us to calculate the fraction of trials, resulting in a smoother switching curve for each different L_Random_ block. After establishing this more reliable switching curve, we were able to determine the trial from the switch for each animal.

### Analysis of behavioral data – reinforcement learning models

The response-by-response behavior of the animal was fitted with eight models: (1) win-stay lose-switch (WSLS); (2) Q-learning (Q-RPE); (3) Q-learning with forgetting (F-Q-RPE); (4) Q-learning with differential forgetting (DF-Q-RPE); (5) F-Q-RPE with choice kernel (F-Q-RPE-CK), which captured the tendency to repeat the same option; (6) DF-Q-RPE with choice kernel (DF-Q-RPE-CK); (7) belief model that uses the prior knowledge of a change point in reward probabilities to make a decision (7) belief model with choice kernel (belief-CK). We will describe these models in detail in the following paragraphs.

For *<u>win-stay lose-switch (WSLS)</u>*, if the last trial was rewarded, the agent would repeat to choose the same option with probability *p*. Else, if the last trial was unrewarded, the agent would switch to choose the other option with probability *p*. This model has 1 free parameter: *p*.

For the three *<u>simple Q-learning models (Q-RPE, F-Q-RPE, DF-Q-RPE)</u>*, the updating rules are as follows. On trial *n*, for a choice *c_n_* that leads to an outcome *r_n_*, the reward prediction error *δ_n_* is:

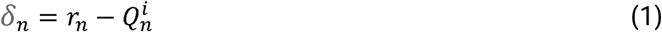

where 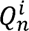 is the action value associated with the chosen action *i*. In our task, there are two options, so *i* ɛ {*L*, *R*}. For the outcome, *r_n_* = 1 for reward, 0 for no reward. The action value for each action is then updated accordingly:

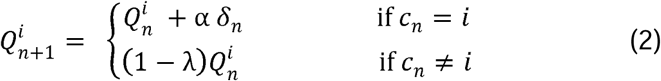

where *α* is the learning rate, *λ* are the forgetting terms for the unchosen action. Then on the next trial, the probability of choosing each action was determined by a softmax rule:

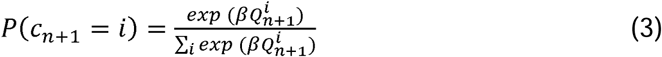

where *β* is the inverse temperature parameter.

The model as stated with 3 free parameters — α, λ, and β — is referred to as Q-learning with differential forgetting (DF-Q-RPE). A special case of this model is when, *λ* = *α*, which is referred to as Q-learning with forgetting (F-Q-RPE). Another special case is when, *λ* = 0, which is referred to as Q-learning (Q-RPE).

For two <u>Q-learning models with choice kernel (*F-Q-RPE-CK, DF-Q-RPE-CK)*</u>, choice kernel was implemented to capture the tendency of choosing the previous choice. We adapted the formulation from (Wilson and Collins, 2019). The choice kernel 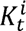 on trial *t* associated with action *i* is updated in a manner analogous to the action values:

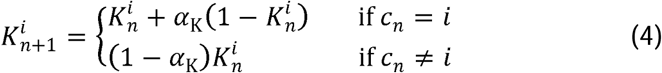

where α_*K*_ is the choice-kernel learning rate. For action selection with both action values and choice kernels, the probability of choosing each action was determined by a softmax rule:

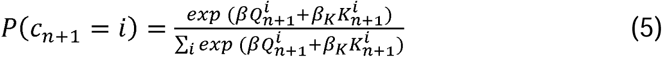

where β and β_*K*_ are the inverse temperature parameters for the action values and choice kernels respectively. Note that the term within the numerator on the right-hand side can be re-arranged:

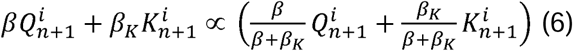

Where (β + β_*K*_) is the effective inverse temperature parameter reflecting the exploration-exploitation balance, and 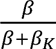 is a ratio indicating the relative reliance on expected reward rather than perseveration in action selection.

### Analysis of behavioral data – belief models

In the <u>belief model</u>, the agent knows two aspects about the task structure. First, the left option can have reward probabilities of either 10% or 70%. It follows that the right option would have the other reward probability. These are the two possible hidden states of the environment. Second, the reward probabilities will reverse with a certain frequency characterized by a hazard rate, *H*. In each trial, the animal has a belief, which consists of the likelihood that the left option has a reward probability of 10%, *ρ_L10_*, and the likelihood that the left option has a reward probability of 70%, *ρ_L70_*. The constraints are that *ρ_L10_* + *ρ_L70_* = 1, *ρ_R10_* =1 - *ρ_L70_* and *ρ_R70_* =1 - *ρ_L70_*. At the start of a session, we set the prior as a uniform distribution, so the *ρ_R10_* = *ρ_L70_* = *ρ_prior_* = 0.5. At the end of each trial, the belief is updated. The possibility of a reward probability switch is considered:

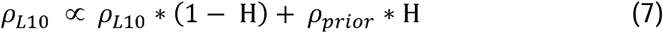

Similarly, *ρ_L70_* is updated and then *ρ_L10_* and *ρ_L70_* are normalized to sum to 1. Next, inference is made based on the outcome following Bayes’ rule, which states that P (belief | observation) = P(belief) * P (observation | belief):

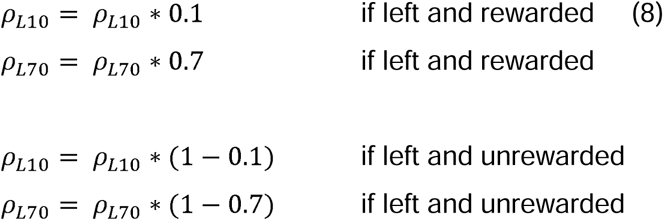

or if the animal chooses right, then *ρ_R10_* and *ρ_R70_* would be updated instead. Again, the probabilities for the belief are normalized to sum to 1. With the updated belief, the expected rewards for the left and right options can be calculated directly, for example:

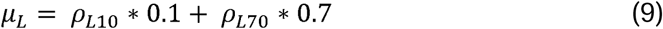

Action selection then proceeds using the same softmax equation as Equations 3 or 5, with the expected rewards replacing the action value terms, for the belief and belief with choice kernels models respectively. The belief model has two free parameters, *H* and *β*. The belief model with choice kernel model has four free parameters *H*, *β*, *α_K_*, and β*_K_*.

### Parameter fitting and model evaluation

For each animal, trials across sessions were concatenated. The values for the free parameters were determined by fitting each model to the concatenated data using the Bayesian adaptive direct search (BADS) algorithm with default settings (Acerbi and Ma, 2017). The initial values for *α, λ, H*, *α_K_*, β and *β_K_* were set to 0.3, 0.3, 0.1, 0.2, 5 and 5 respectively. The lower bound of parameters were set to 0 and the upper bound was set to 100 for inverse temperatures and 1 for the rest of the parameters. To evaluate the models, we calculated the Bayes information criterion (BIC).

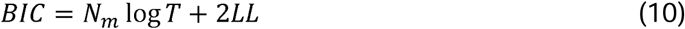

where N_m_ is the number of parameters in model m. T is the number of trials used to estimate the parameters and LL is the negative log-likelihood value at the best fitting parameter settings. The model that best fits the data should have the smallest BIC score as the positive effect of the number of parameters, N_m_, has an explicit penalty for free parameters.

The parameters used for the simulated data for the belief model with choice kernel was the best-fitting parameters of one animal (H = 0.320, *β* = 1.387, *α_k_*=0.468, *β_K_* = 2.543 with 300,000 trials, approximately 9000 switches as in the experiment data). The belief model with choice kernel was used to analyze the latent variables for lesion and photostimulation data.

To fit the lesion data, we modified the belief-CK model. Different parameters for hazard rate and choice kernel learning rate were used depending on if the animal’s choice in the current trial is ipsilateral or contralateral to the lesion side. This yields an expanded model with 6 parameters: *H_lesion_, H_Contra_, α_K lesion_, α_K Control_, β*, and *β_K_*. To fit the data on a per-animal basis, trials across sessions before lesion were concatenated, and trials across sessions after lesion were concatenated. To fit the data on a per-session basis, we estimated the parameters for each session.

To fit the optogenetics data, we modified the belief-CK model. Different parameters for hazard rate and choice kernel learning rate were used depending on if the animal’s choice in the current trial is ipsilateral or contralateral to the photostimulated side, or if the animal’s choice occurred in a control trial with no photostimulation. Different parameters for inverse temperatures were used depending on if the current trial was photostimulation or control. This yields an expanded model with 10 parameters: *H_ipsi stim_, H_Contra stim_, H_Control_, α_K ipsi stim_, α_K Contra stim_, α_K Control_, β_Control_, β_stim_, β_K Control_*, and *β_K stim_*. Per each animal fittings, trials across sessions for pre-choice stimulation and post-choice stimulation were concatenated. Per session fittings was not used for this dataset as total number of trials within a session did not give a reliable estimate for the ten parameters fittings.

### Statistical Analyses

All statistical analyses were completed using MATLAB (version 2019b, MathWorks). Three-way ANOVA was used to examine the effect of the lesion on behavioral performance. For datasets with a matched number of data points, Wilcoxon signed-rank test was used; otherwise, Wilcoxon ranked sum test was used. Unless otherwise specified, we used an alpha level of 0.05 for all statistical tests.

## Acknowledgments

We thank Neil Savalia and John-Anthony Fraga for assistance with behavioral training, Michael Siniscalchi for use of help with the behavioral apparatus, Farhan Ali for advice on histology, Lucas Pinto, Stephan Thiberge and David Tank for sharing design of photostimulation rig, and Alireza Soltani for comments on an earlier version of the analysis. This work was supported by NIH/NIMH grants R01MH112750 (A.C.K.), R21MH118596 (A.C.K.), F32NS101871 (L.P.), K99MH120047 (L.P.), China Scholarship Council-Yale World Scholars Fellowship (H.W.), Gruber Science Fellowship (H.K.O.), NIH training grant T32NS007224 (H.K.O.), and Kavli Institute for Neuroscience Postdoctoral Fellowship (H.A.).

## Author Contributions

H.A. and A.C.K. conceived the project. H.A. performed all surgeries. H.A. and C.E.M performed behavioral training including lesion and optogenetic inactivation experiments, and histology. H.K.O. performed the ALM inactivation experiment. H.W., H.A., and A.C.K. put together the optogenetic activation rig, based on L.P.’s plans. H.A. and A.C.K. analyzed the data and wrote the paper with input from all other authors.

## Supplementary Information

**Supplementary Figure 1.1:**
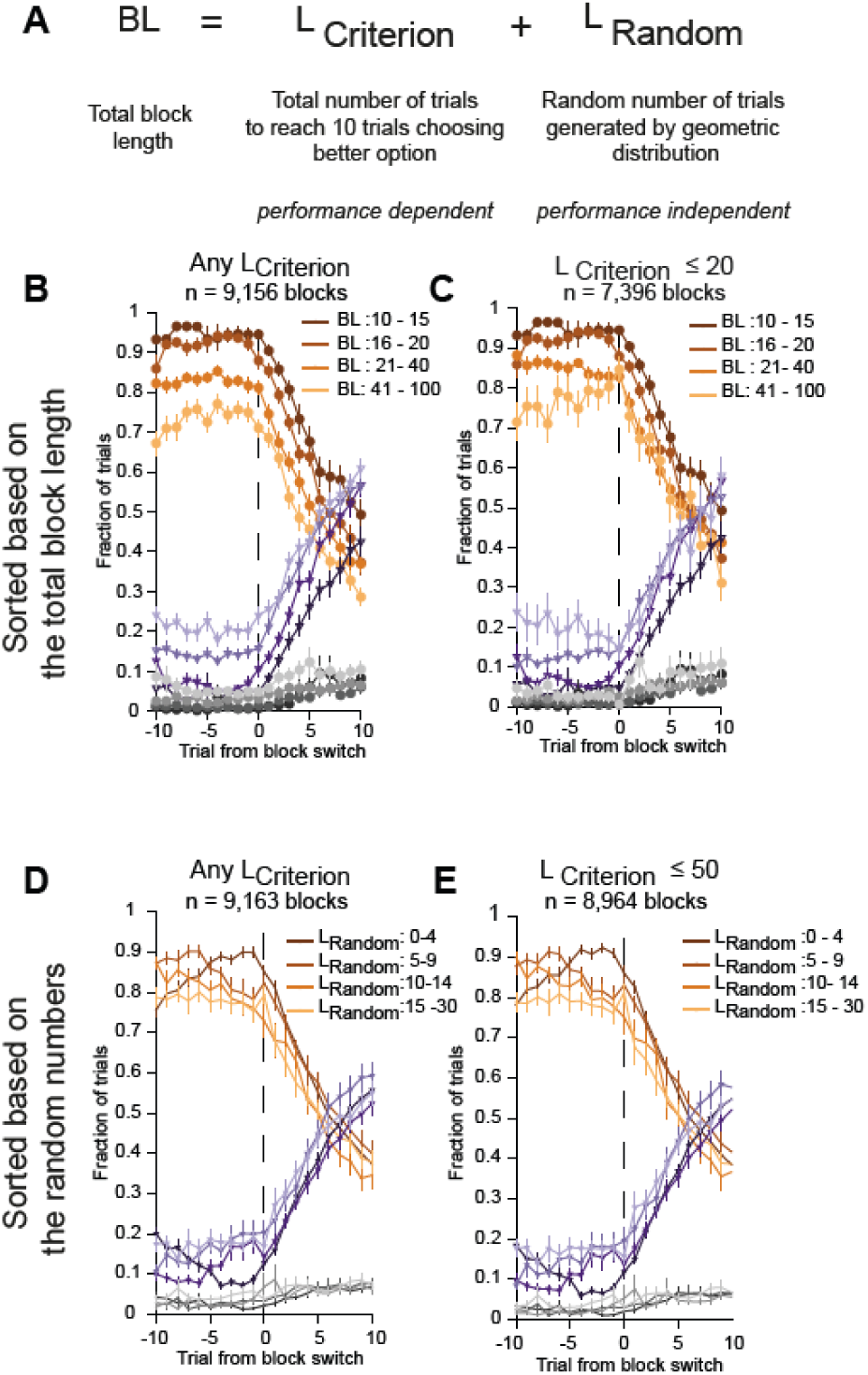
Animal’s choice behavior around block switches with different performance criteria. (A) The equation for the switching condition, or block length (BL), which is the sum of L_criterion_ and L_Random_. (B) Choice behavior around block switches, plotted separately for 4 ranges of BL. Mean values and SEM for all animals. All data were included. (C) Similar to (B), including data in which L_criterion_ ≤ 20 trials. (D) Choice behavior around block switches, plotted separately for 4 ranges of L_Random_. Mean values and SEM for all animals. All data were included. (E) Similar to (C), including data in which L_criterion_ ≤ 50 trials.

**Supplementary Figure 1.2:**
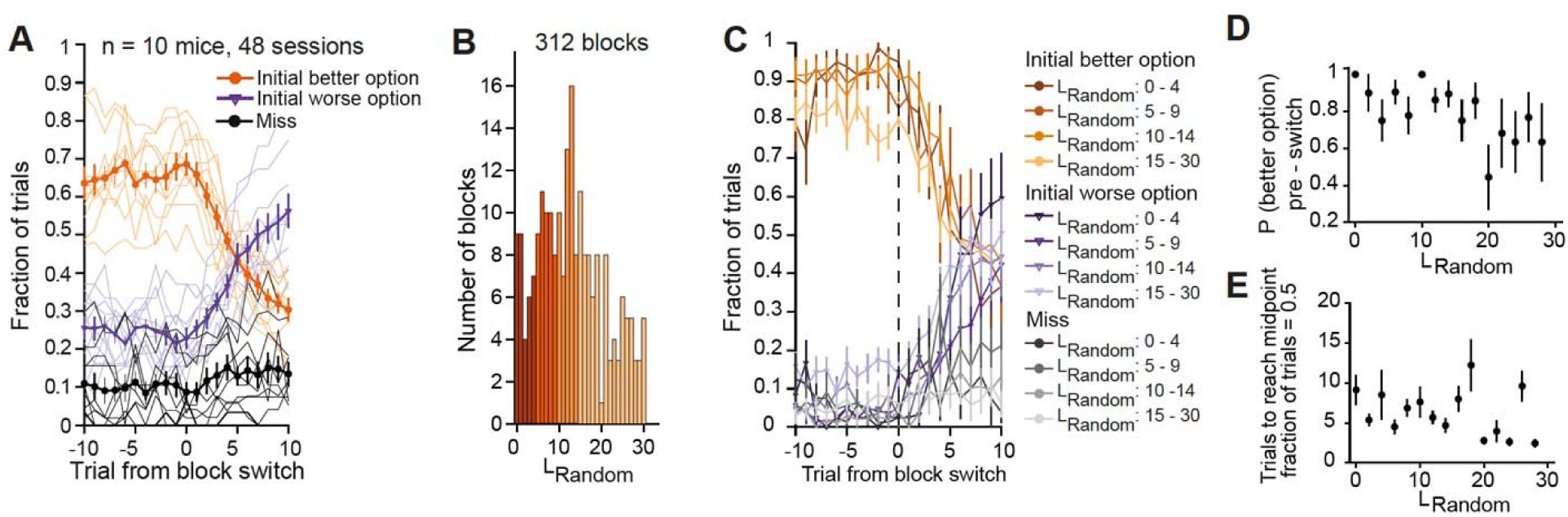
Animal’s choice behavior in a task variant without L_criterion_. (A) Choice behavior around block switches in a task variant without L_criterion_. All the trial timing and reward probabilities are identical, except the switching condition consists of only L_Random_. Thin line, mean values for individual animal. Thick line, mean values and SEM for all animals. (B) Histogram of L_Random._ for all blocks. Colors indicate the 4 ranges of L_Random_ for subsequent analyses. (C) Choice behavior around block switches, plotted separately for the 4 ranges of L_Random_. Mean values and SEM for all animals. (D) The probability of choosing the better option on the trial immediately preceding the switch, as a function of L_Random_ for the block preceding the switch (in 2 datapoints bin; main effect of L_Random_: F (14, 204) = 1.8068, *P* = 0.0394 one-way ANOVA) Mean values and SEM for all animals. (E) The number of trials to reach midpoint (when animal is equally likely to choose either option) as a function of L_Random_ for the block preceding the switch (in 2 datapoints bin; main effect of L_Random_: F (14, 120) = 1.0734, *P* = 0.3883 one-way ANOVA). Mean values and SEM for all animals. n = 10 mice, 48 sessions, 312 blocks.

**Supplementary Figure 2.1:**
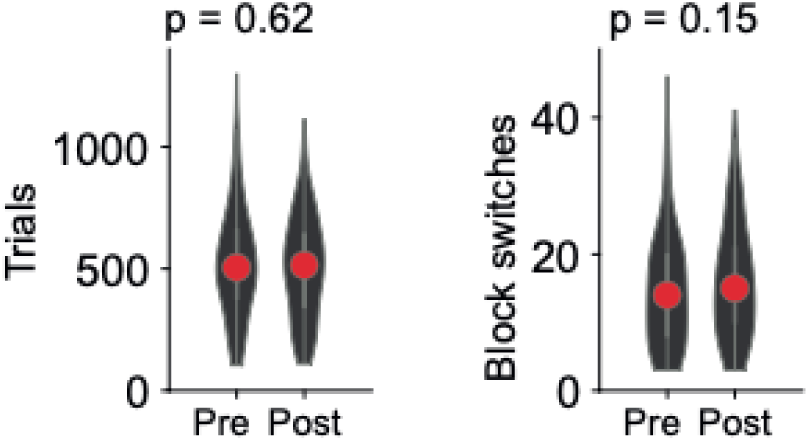
No decrease in overall performance after unilateral lesion of ACAd/Mos. The total number of trials and block switches per session before (pre) and after (post) the unilateral lesion.

**Supplementary Figure 2.2:**
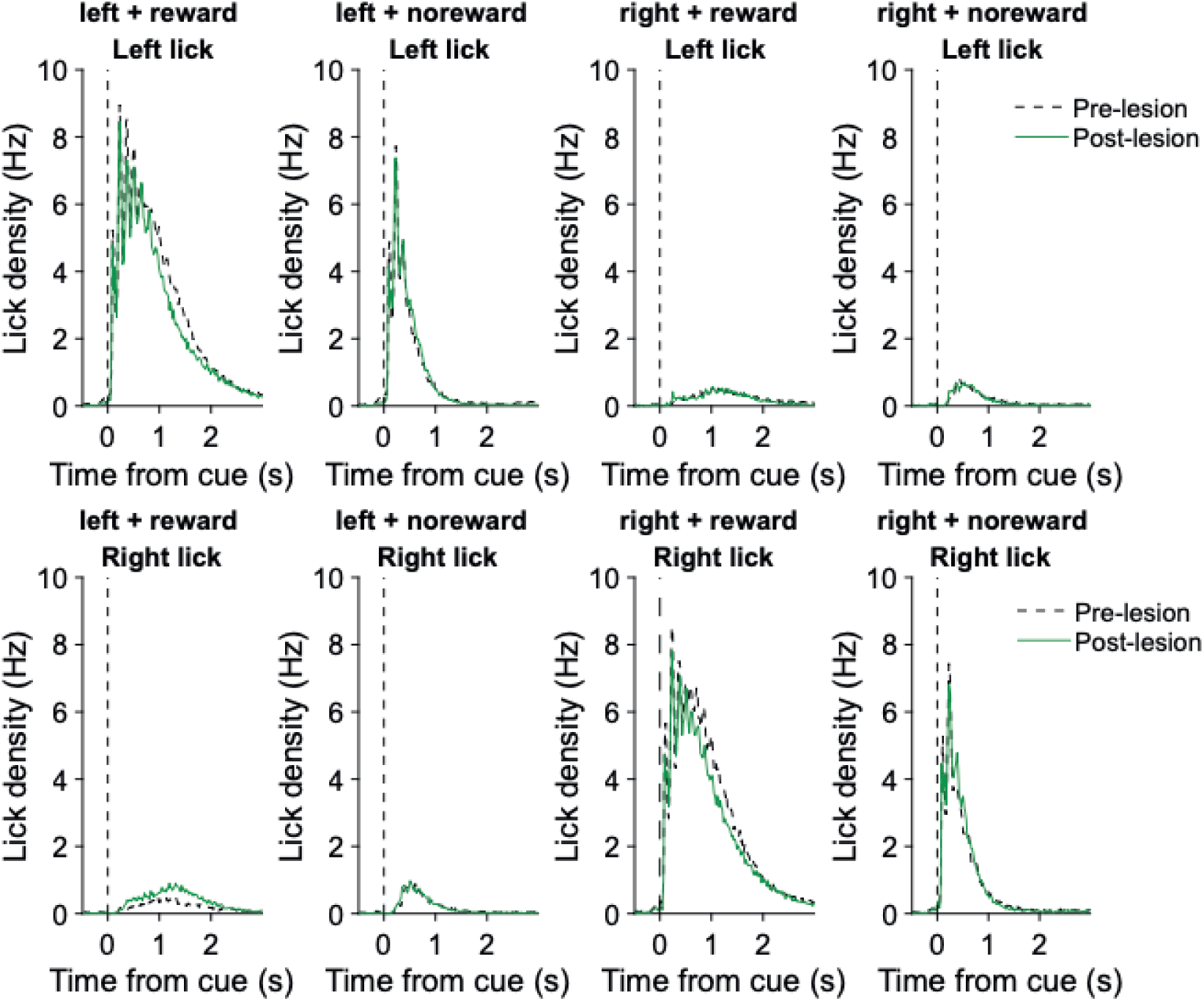
No motor deficits after unilateral lesion of ACAd/Mos. Mean left and right lick density for each possible combination for choice (left or right) and outco e (reward or no reward). No significant difference was detected between pre- and post-unilateral lesion.

**Supplementary Figure 3.1:**
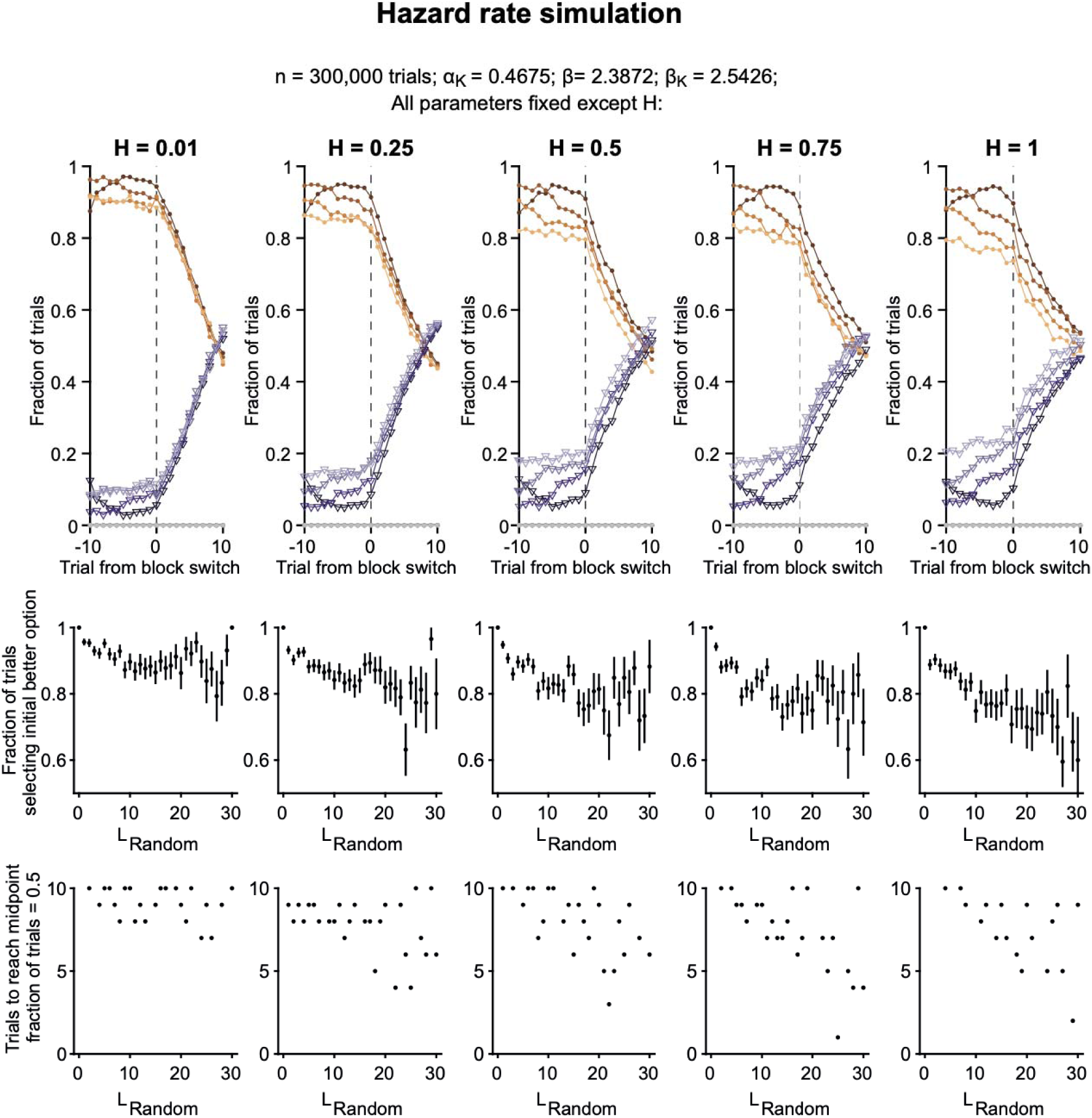
Belief-CK model: effect of varying the hazard rate. The belief-CK model was used to simulate an agent’s choice behavior in the two-armed bandit task with probabilistic reward reversal. Parameters were selected based on the best fitting values from an animal. Each column shows the results using a different hazard rate (= 0.01, 0.25, 0.5, 0.75, 1) while all other parameters were kept constant (n = 300,000 trials, = 1.387, =0.468, = 2.543). Top row shows the mean fraction of trials choosing the better and worse options for 4 different L_Random_ ranges for 10 trials before and after the block switch. Middle row shows the P (better option) _pre-switch_ as a function of L_Random_. Mean and SEM. Bottom row shows the mean number of trials to reach midpoint as a function of L_Random_.

**Supplementary Figure 3.2:**
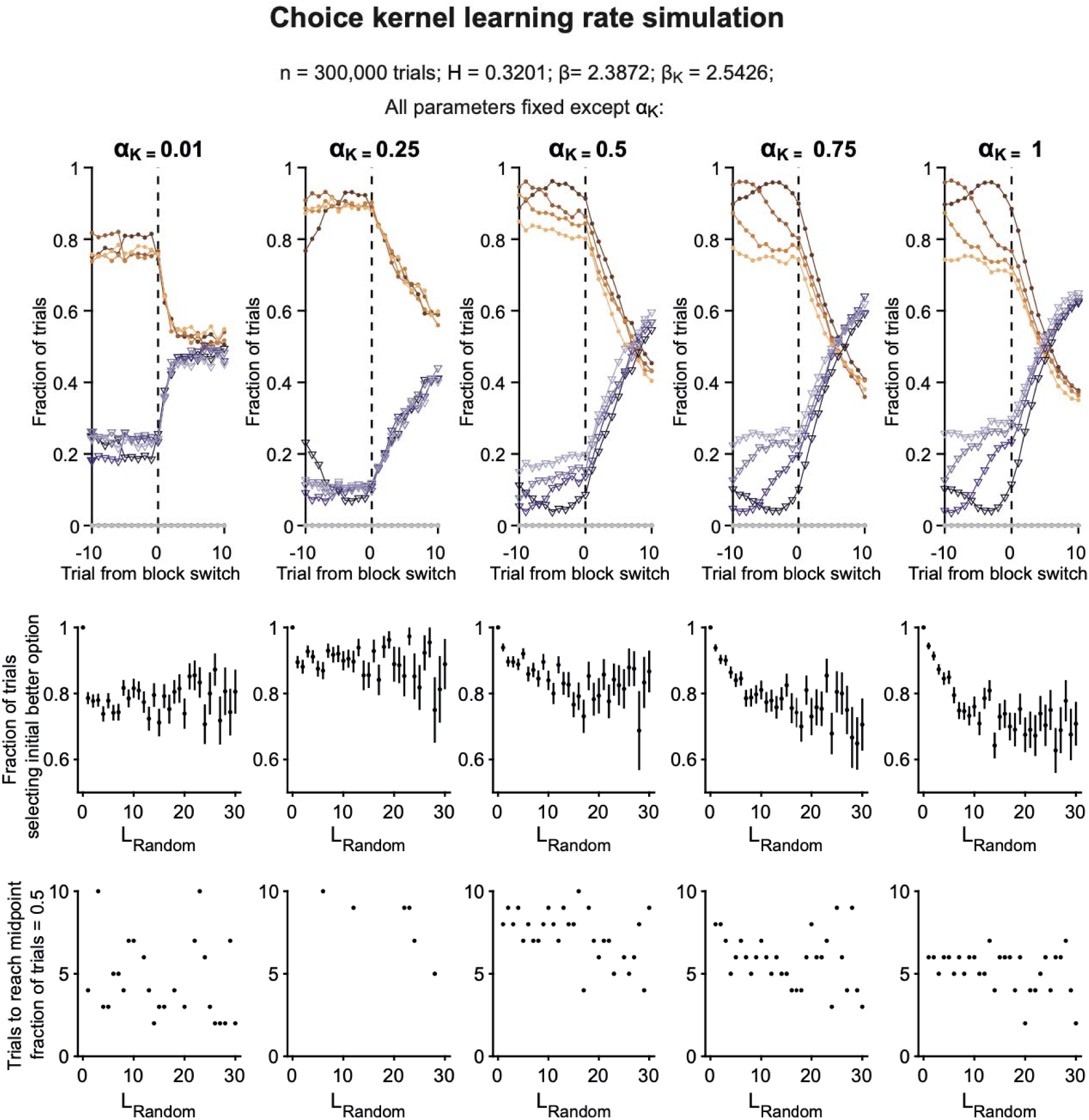
Belief-CK model: effect of varying choice kernel learning rate. Similar to Supplementary Figure 3.1, with different choice kernel learning rates (= 0.01, 0.25, 0.5, 0.75, 1) while all other parameters were kept constant (n = 300,000 trials, = 0.320, = 1.387, = 2.543).

**Supplementary Figure 3.3:**
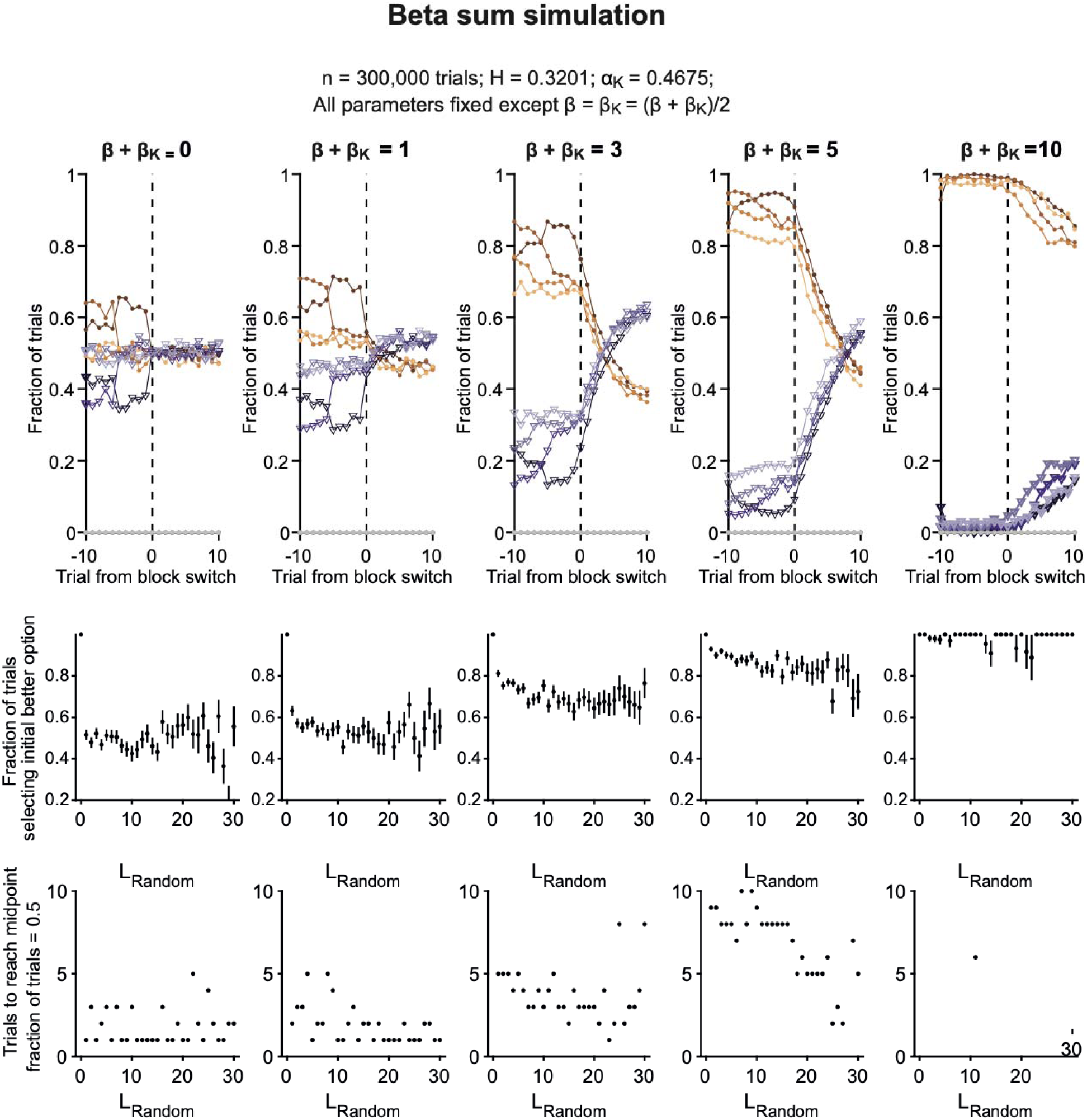
Belief-CK model: effect of varying beta sum. Similar to Supplementary Figure 3.1, with different beta sum (= 0, 1, 3, 5, 10) while all other parameters were kept constant (n = 300,000 trials, = 0.320, = 0.468). was set to equal to.

**Supplementary Figure 3.4:**
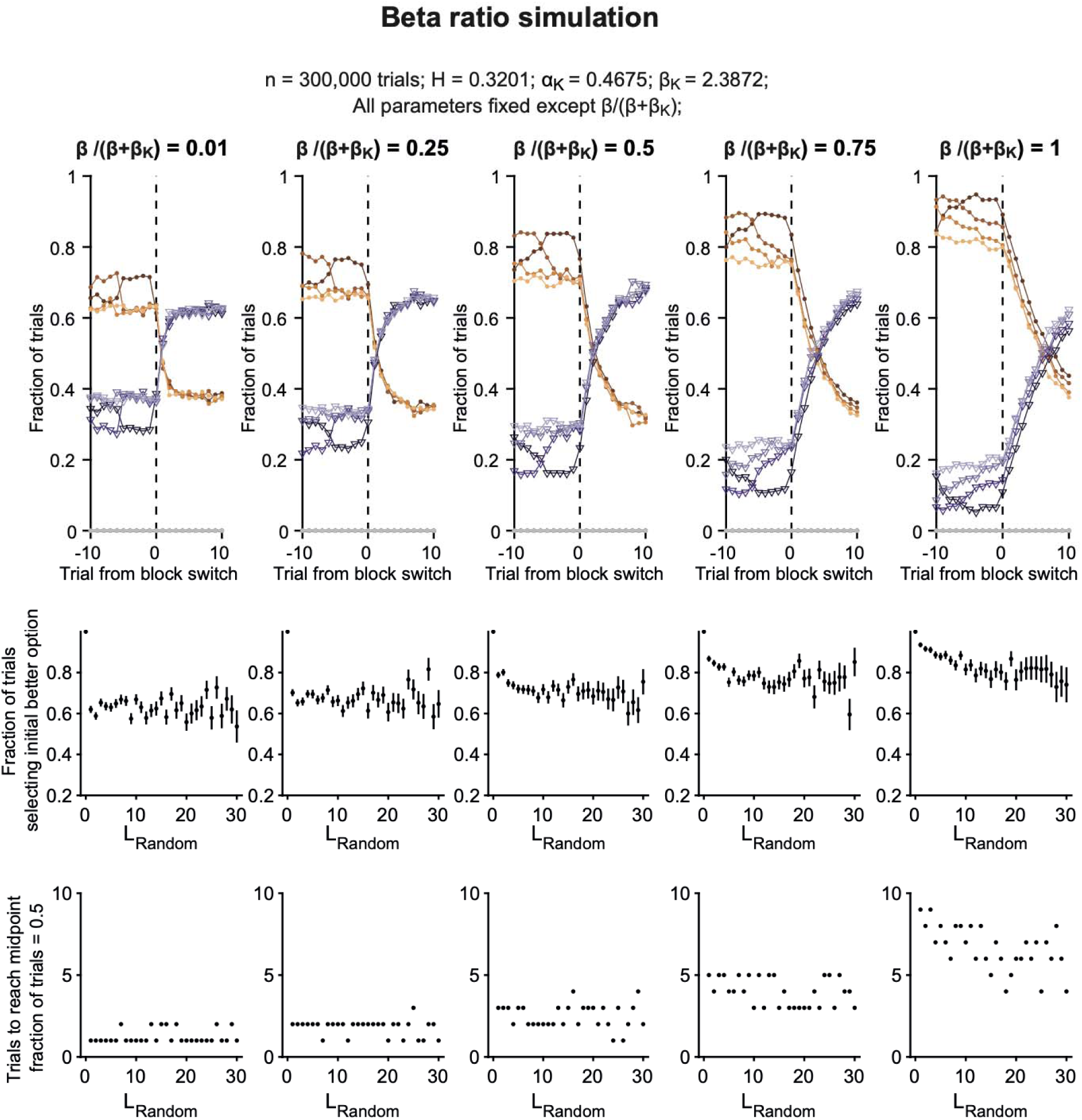
Belief-CK model: effect of varying beta ratio. Similar to Supplementary Figure 3.1, with different beta ratios (0.01, 0.25, 0.5, 0.75, 1) while all other parameters were kept constant (n = 300,000 trials, = 0.320, =0.468, = 2.543). was fixed and was calculated based on the beta ratio values.

**Supplementary Figure 3.5:**
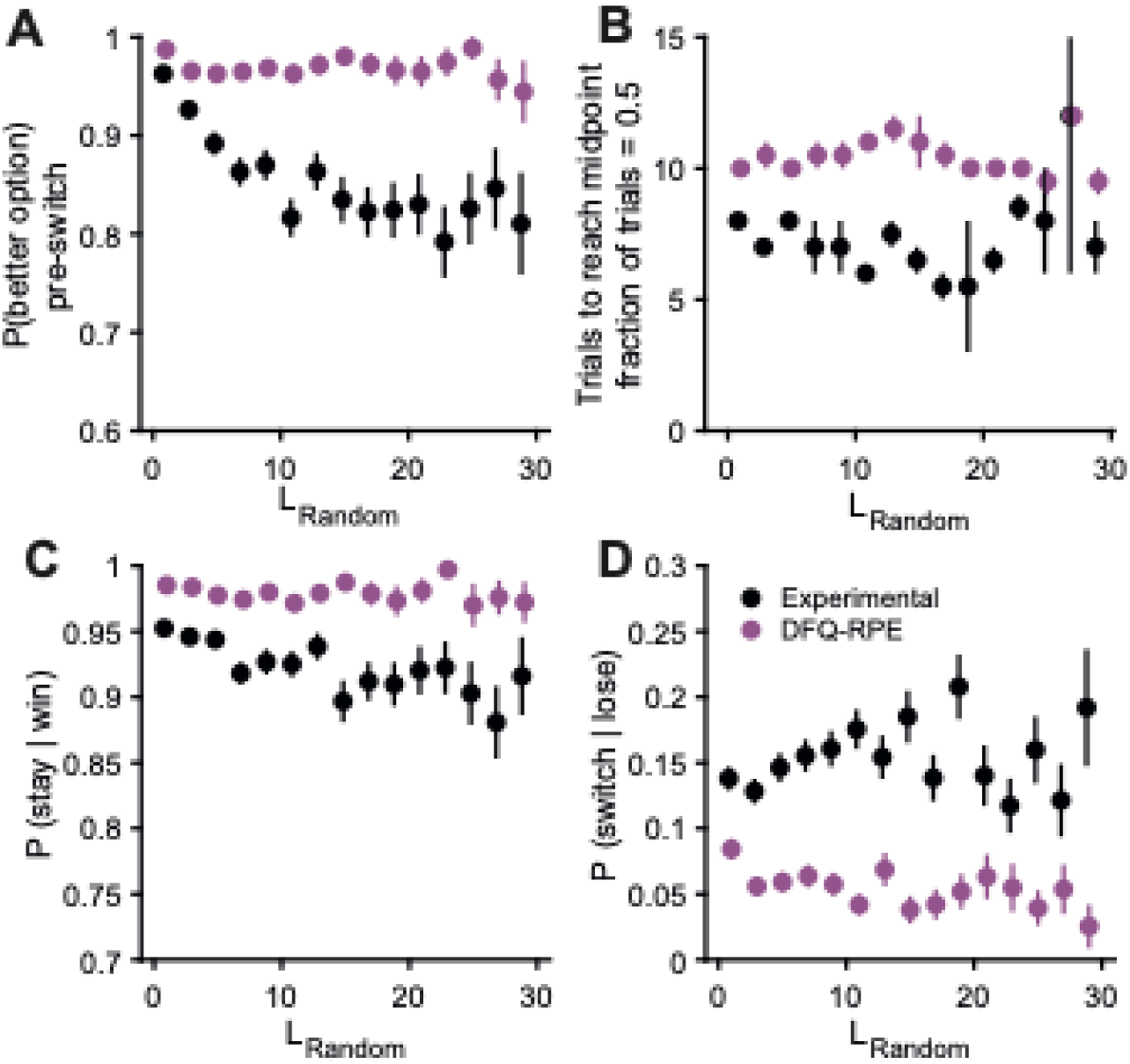
DF-Q-RPE algorithm cannot reproduce the L_Random_-dependent trends in the experimental data. **(A)** The probability of choosing the better option on the trial immediately preceding the switch, as a function of L_Random_ for the block preceding the switch. Black, mice. Purple, simulated performance using the DF-Q-RPE model with best-fitting parameters. Mean values and SEM for all animals. **(B)** Similar to (A) for number of trials to reach midpoint (when animal is equally likely to choose either option). **(C)** Similar to (A) for the tendency to win-stay on the 5 trials preceding the switch. **(D)** Similar to (A) for the tendency to lose-switch on the 5 trials preceding the switch. Mean and SEM. n = 31 mice, 617 sessions.

**Supplementary Figure 5.1:**
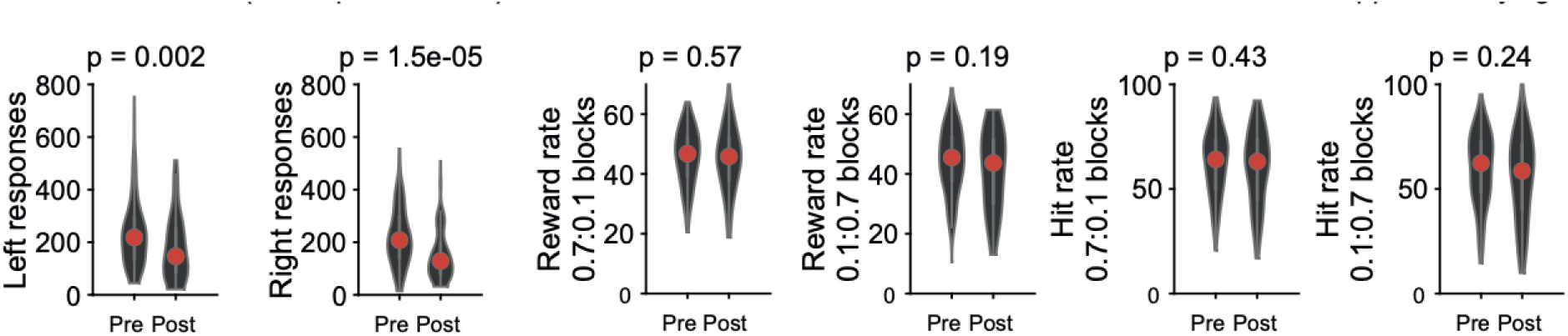
Fewer trials but similar performance after bilateral lesion of ACAd/Mos. The total number of left- and right-responding trials, reward rates, and hit rates before (pre) and after (post) the lesion.

**Supplementary Figure 5.2:**
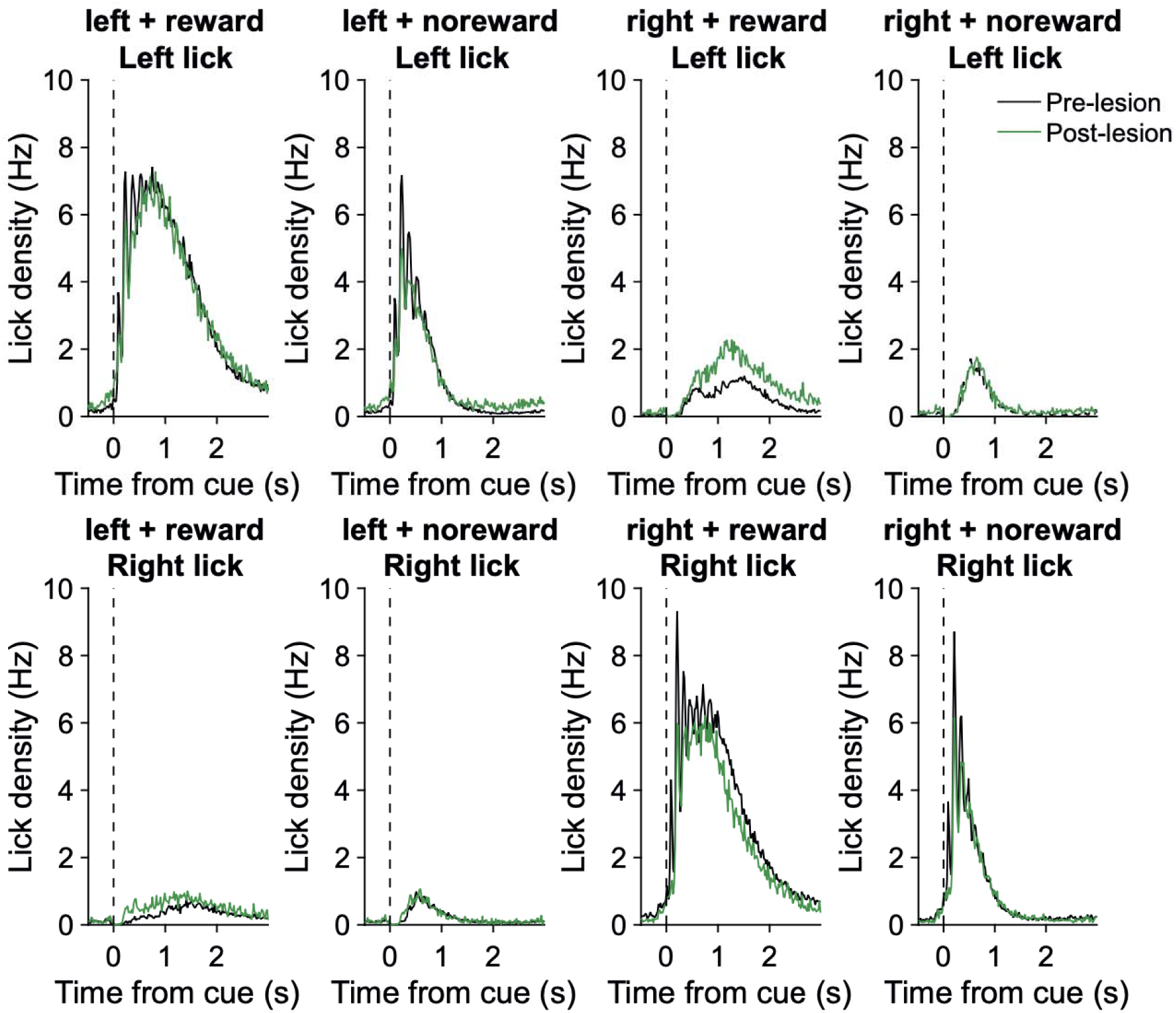
No motor deficits after bilateral lesion of ACAd/Mos. Mean left and right lick density for each possible combination for choice (left or right) and outco e (reward or no reward). No significant difference was detected between pre- and post-bilateral lesion.

**Supplementary Figure 6.1:**
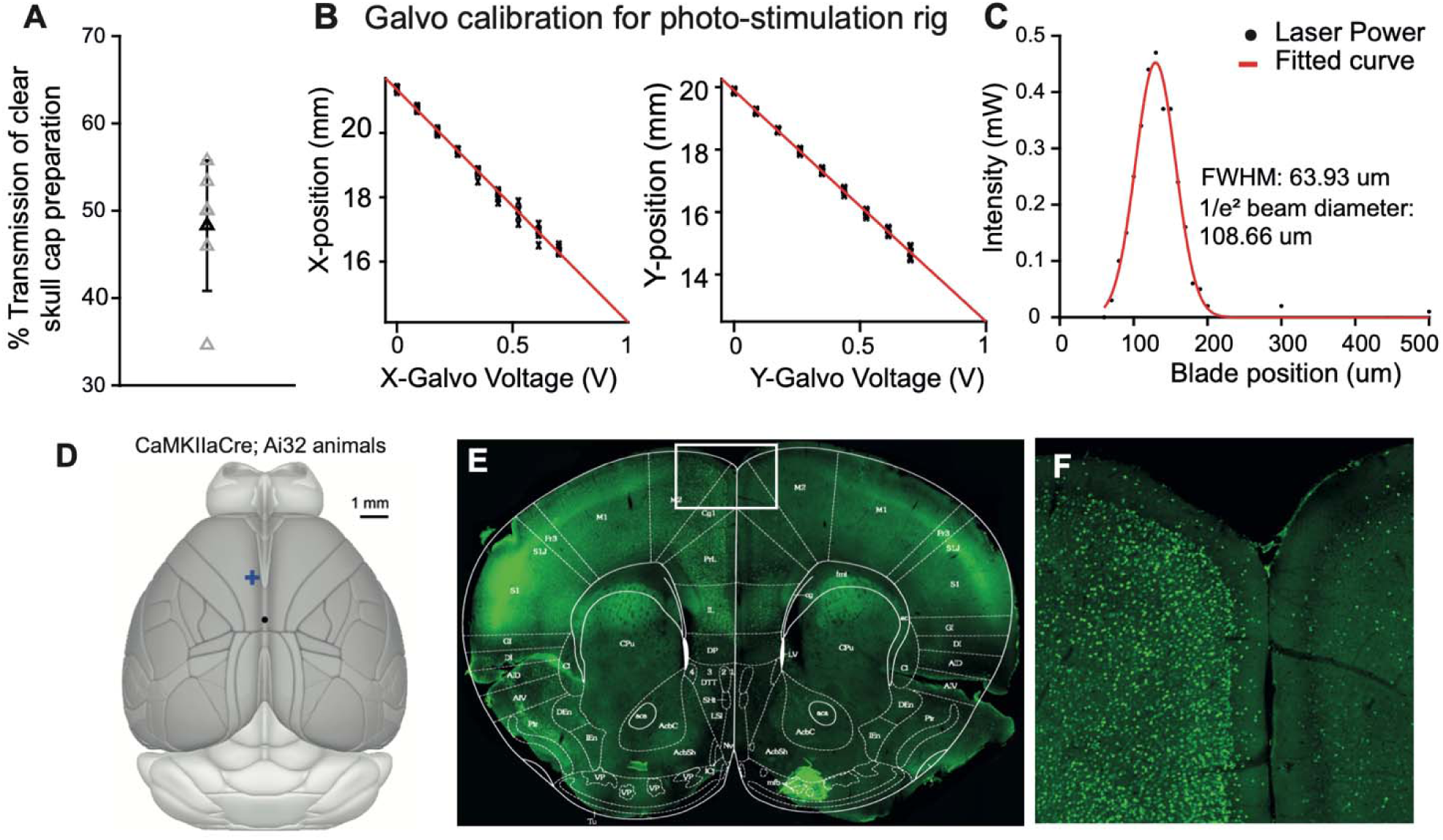
Validation of the laser steering system for optogenetic manipulation: characterization and c-Fos staining. (A) Optical transmission of the clear skull cap preparation was measured by illuminating with a laser and recording intensity using a power meter. Mean and SEM. n = 5. (B) Linearity of the galvanometers in the x and y directions. (C) Beam profile was measured at the sample plane by inserting and moving a razor blade across the plane using a micromanipulator. (D - F) In *CaMKIIa^Cre^;Ai32* animals, cortical excitatory neurons express ChR2. After unilateral photostimulation of the left ACAd/MOs region (40 Hz, 1.5 mW, 1 min on then 1 min off repeatedly for 20 min), immunohistostaining with a c-Fos antibody showed elevated signals.

**Supplementary Figure 6.2:**
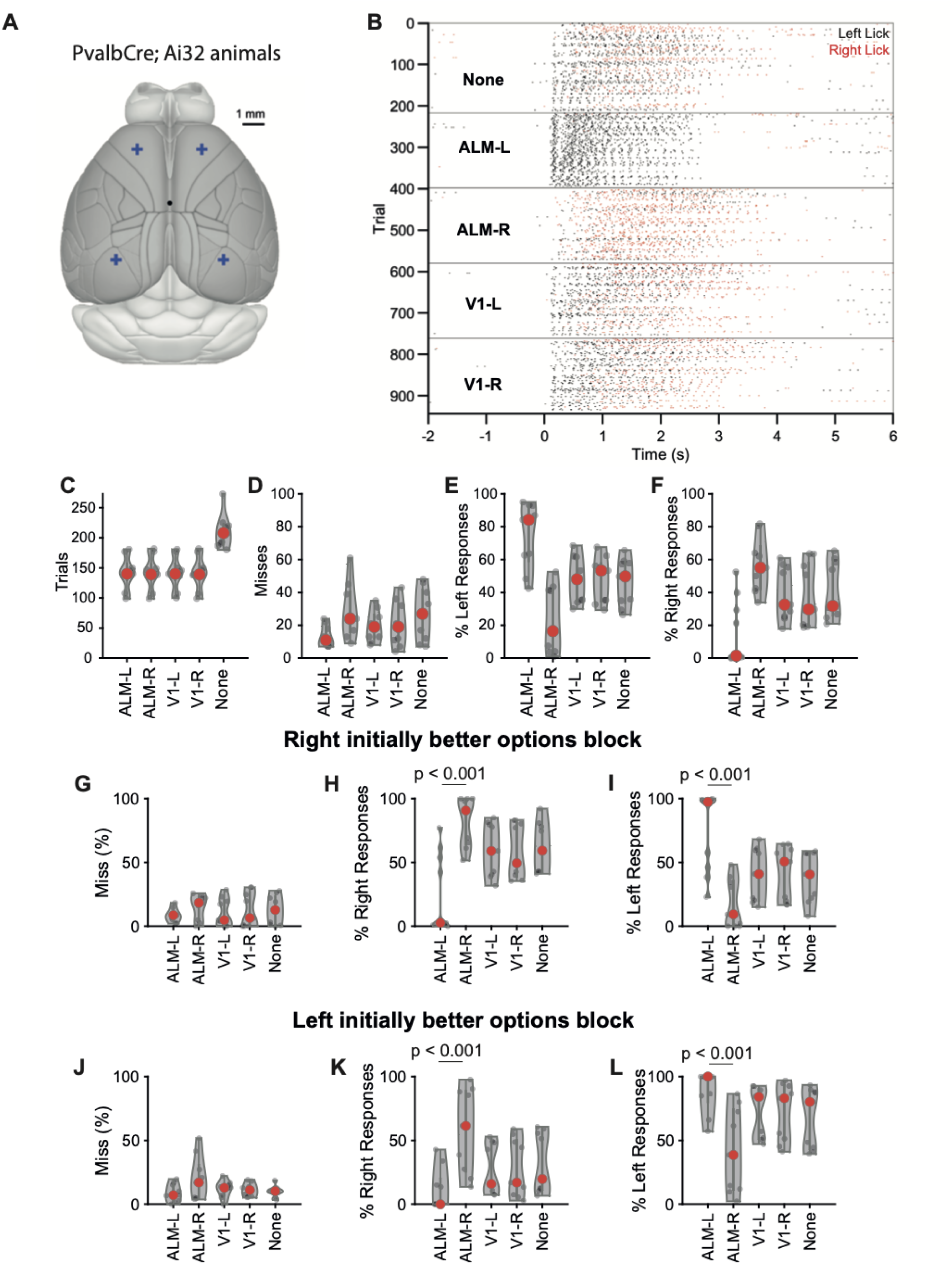
Inactivating left and right ALM during two-armed bandit task. **(A)** In *Pvalb^Cre^;Ai32* animals, parvalbumin-expressing neurons including fast-spiking interneurons in the neocortex express ChR2. Photostimulation of a brain region drives spiking in the interneurons, which in turn suppresses excitatory activity. Lick raster recorded in an example session, in which trials were sorted based on the photostimulation (None: no stimulation; ALM-L: left anterior lateral motor cortex, AP=2.5 mm, ML=-1.5 mm; ALM-R: right anterior lateral motor cortex, AP=2.5 mm, ML=1.5 mm; V1-L: left primary visual cortex, AP=-2.7 mm, ML=-2.5 mm; V1-R: right primary visual cortex, AP=-2.7 mm, ML=2.5 mm). **(B)** The number of trials of each type per session. **(C)** Percent of trials resulted in a miss, as a function of trial type. **(D)** Percent of trials resulted in a left response, as a function of trial type. **(E)** Percent of trials resulted in a right response, as a function of trial type. These results show that transient inactivation of ALM increased ipsilateral responses at the expense of contralateral responses. 9 sessions from 3 animals.

**Supplementary Table 2.1:** The results of three-way between-subjects ANOVA with factors of lesion (pre- and post-lesion), side (lesion blocks and Contra blocks), and L_Random_ (4 L_Random_ ranges) for P(better option)_pre-switch_, trials to reach midpoint and hit rates. p < 0.05 in bold. All dependent variables calculated for each block across sessions. (Error = 3285; 2399; 3285; 3285; 2816;3187 for P (better option) _pre-switch_, trials to reach midpoint and hit rates respectively

|  |  | P(better option)<br>pre-switch |  | Trials to reach<br>midpoint |  | Hit rates |  |
| --- | --- | --- | --- | --- | --- | --- | --- |
|  | df | F | p | F | p | F | p |
| *Lesion* | 1 | **32.71** | **.000** | **33.36** | **.000** | **4.87** | **.027** |
| *Side* | 1 | 0.63 | .857 | **8.68** | **.003** | **4.02** | **.045** |
| *L<sub>Random</sub>* | 3 | **57.67** | **.000** | 0.52 | .666 | **21.46** | **.000** |
| *Lesion\*Side* | 1 | 1.42 | .233 | **4.30** | **.038** | 0.86 | .355 |
| *Lesion\*L<sub>Random</sub>* | 3 | **2.71** | **.044** | 0.57 | .635 | **3.06** | **.027** |
| *L<sub>Random</sub>\*Side* | 3 | 0.18 | .910 | **4.30** | **.004** | 0.49 | .688 |
| *Lesion\*L<sub>Random</sub>\*Side* | 3 | 2.40 | .066 | 1.60 | .187 | 1.07 | .361 |

**Supplementary Table 5.1:** The results of two-way between-subjects ANOVA for bilaterally injected animals with factors of lesion (pre- and post-lesion and L_Random_ (4 L_Random_ ranges) for P(better option) _pre-switch_, trials to reach midpoint and hit rates. p < 0.05 in bold. All dependent variables calculated for each block across sessions. (Error = 1356;911;1356; for P (better option) _pre-switch_, trials to reach midpoint and hit rates respectively

|  |  | P(better option)<br>pre-switch |  | Trials to reach<br>midpoint |  | Hit rates |  |
| --- | --- | --- | --- | --- | --- | --- | --- |
|  | df | F | p | F | p | F | p |
| *Lesion* | 1 | 1.968 | .161 | 0.001 | .974 | 1.0814 | 0.2986 |
| *L<sub>Random</sub>* | 3 | 20.179 | .000 | 1.525 | .206 | 1.8643 | 0.1337 |
| *Lesion\*L<sub>Random</sub>* | 3 | 1.285 | .278 | 0.202 | .895 | 0.4492 | 0.7179 |

**Supplementary Table 5.2:** The results of two-way between-subjects ANOVA for saline injected animals with factors of lesion (pre- and post-lesion and L_Random_ (4 L_Random_ ranges) for P (better option) _pre-switch_, trials to reach midpoint, hit rates, P(lose | switch), P(win | stay). p < 0.05 in bold. All dependent variables calculated for each block across sessions. (Error = 1871;1217; 1871 for P (better option) _pre-switch_, trials to reach midpoint and hit rates respectively.

|  |  | P(better option)<br>pre-switch |  | Trials to reach<br>midpoint |  | Hit rates |  |
| --- | --- | --- | --- | --- | --- | --- | --- |
|  | df | F | p | F | p | F | p |
| *Lesion* | 1 | 0.0167 | .897 | 9.7782 | .002 | 0.0325 | 0.857 |
| *L<sub>Random</sub>* | 3 | 20.430 | .000 | 1.0813 | .356 | 9.624 | 0.000 |
| *Lesion\*L<sub>Random</sub>* | 3 | 0.082 | 0.969 | 0.0594 | .981 | 0.190 | 0.903 |

